# Monitoring rapid evolution of plant populations at scale with Pool-Sequencing

**DOI:** 10.1101/2022.02.02.477408

**Authors:** Lucas Czech, Yunru Peng, Jeffrey P. Spence, Patricia L.M. Lang, Tatiana Bellagio, Julia Hildebrandt, Katrin Fritschi, Rebecca Schwab, Beth A. Rowan, GrENE-net consortium, Detlef Weigel, J.F. Scheepens, François Vasseur, Moises Exposito-Alonso

## Abstract

The change in allele frequencies within a population over time represents a fundamental process of evolution. By monitoring allele frequencies, we can analyze the effects of natural selection and genetic drift on populations. To efficiently track time-resolved genetic change, large experimental or wild populations can be sequenced as pools of individuals sampled over time using high-throughput genome sequencing (called the Evolve & Resequence approach, E&R). Here, we present a set of experiments using hundreds of natural genotypes of the model plant *Arabidopsis thaliana* to showcase the power of this approach to study rapid evolution at large scale. First, we validate that sequencing DNA directly extracted from pools of flowers from multiple plants -- organs that are relatively consistent in size and easy to sample -- produces comparable results to other, more expensive state-of-the-art approaches such as sampling and sequencing of individual leaves. Sequencing pools of flowers from 25-50 individuals at ∼40X coverage recovers genome-wide frequencies in diverse populations with accuracy *r* > 0.95. Secondly, to enable analyses of evolutionary adaptation using E&R approaches of plants in highly replicated environments, we provide open source tools that streamline sequencing data curation and calculate various population genetic statistics two orders of magnitude faster than current software. To directly demonstrate the usefulness of our method, we conducted a two-year outdoor evolution experiment with *A. thaliana* to show signals of rapid evolution in multiple genomic regions. We demonstrate how these laboratory and computational Pool-seq-based methods can be scaled to study hundreds of populations across many climates.

## Introduction

How fast a species can adapt to different environments from standing within-species genetic variation is a burning question in evolutionary ecology and genetics. A powerful approach to study environment-driven adaptation is provided by field experiments in which multiple genotypes of a species are grown together and traits and fitness are measured (Clausen et al., 1941; Kingsolver et al., 2001; Savolainen et al., 2013). Such experiments, typically conducted within a single generation, have allowed measuring the strength of natural selection over phenotypic traits or genetic variants, which is often strong (Exposito-Alonso et al., 2019; Kingsolver et al., 2001; Siepielski et al., 2017; Thurman and Barrett, 2016). Such studies often cannot measure the response to selection—evolutionary change—of a population, as this depends on the genetic trait architecture (Bergland et al., 2014; Walsh and Blows, 2009) and environmental fluctuation over time (Bergland et al., 2014), which has led to inconsistent long-term trait changes in populations (Merilä et al., 2001). Highly-replicated multi-year experiments where phenotypes and genomic variation are tracked would be ideal to study these evolutionary forces and robustly test the predictability of evolution (Grant and Grant, 2002; Nosil et al., 2018) .

An opportunity to conduct multi-generational experiments to study evolution over time is the so-called “Evolve & Resequence” (E&R) approach (Schlötterer et al., 2015; Turner et al., 2011). E&R experiments leverage cost-effective, high-throughput sequencing to study the frequency of genome-wide variants or genotypes of a population over time, especially when different populations are subject to different environmental conditions that may reveal phenotypic variation (Bastide et al., 2013; Schlötterer et al., 2015). Such frequency trajectories capture evolutionary forces such as drift and natural selection in action. This approach has been inspired in early experiments in bacterial and animal model systems such as *Escherichia coli* and *Drosophila melanogaster* (Bergland et al., 2014; Good et al., 2017; Schlötterer et al., 2014). In the traditional genome sequencing approach, each individual is processed independently into one DNA sequencing library. The most common sequencing approach for E&R is Pool-Sequencing, where multiple individuals sampled from the same population are processed into a single DNA sequencing library (Futschik and Schlötterer, 2010). While individual haplotypes are lost in the Pool-Seq approach, population-level allele frequencies are obtained in a cost-effective manner (Schlötterer et al., 2014). The Pool-Seq approach has been typically applied on a single population over time to study rapid selective sweeps (Iranmehr et al., 2017) and quantitative trait evolution (Endler et al., 2016). Parallel E&R experiments across large environmental gradients could enable the study of population (mal)adaptation across present climates and inform future responses (Capblancq et al., 2020). Combining Pool-Seq experiments, which subject the same starting genetic variation to an environmental condition, with landscape genomic approaches that aim to detect climate-driven natural selection or sweeps in the presence of population confounders (Günther and Coop, 2013; Hancock et al., 2011; Pfenninger et al., 2021), could be a powerful approach to depict how climate impacts evolutionary genetic processes leading to adaptation and extinction.

To enable globally-distributed E&R experiments to study climate adaptation, two key innovations are necessary beyond lowering sequencing costs, which we address here: (1) making library preparation scalable to thousands of whole-genome samples, and (2) standardizing computational genomics software that allow researchers to analyze thousands of population samples, akin in speed to single-genome data structures and libraries such as HTSlib (Bonfield et al., 2021). To achieve the first goal, we hereby document Pool-seq protocols to reduce the preparation time (to ∼2 h/96 pooled samples) and cost (to ∼$3/pooled sample) for genomic DNA library preparation using Tn5 transposase. These were adapted from the Baym (2015) and Rowan studies (2015), and were tested here for Pool-Sequencing approaches. For the second goal, we developed a new C++ implementation for fast computing of population genetic statistics for Pool-Seq, `g`_r_`enedalf`, (Czech and Exposito-Alonso, 2022), reimplementing the equations of the original Perl-based PoPoolation software (Kofler et al., 2011a, 2011b). Our implementation now offers ∼100-fold speed improvements, allowing analyses of thousands of pooled libraries in minutes rather than days (Czech and Exposito-Alonso, 2022). These methods can be applied to any organisms, and we demonstrate the utility and power of the approach with the diploid annual plant *Arabidopsis thaliana*. We showcase our methods’ efficacy for studying rapid adaptation in the context of plant evolutionary ecology, a field that typically uses individual-based methods such as common garden experiments and within-generation fitness assays to understand natural selection in different environments (Anderson and Wadgymar, 2019; Brachi et al., 2010; Exposito-Alonso et al., 2019; Fournier-Level et al., 2011; Lovell et al., 2021; Lowry et al., 2009; Monnahan et al., 2020) .

In this article, we describe our E&R design, Pool-Seq protocols, and computational approaches for a set of four experiments using natural genotypes of *A. thaliana*. We provide evidence that our simple and affordable large-scale experimental setup can generate allele frequency data with quality comparable to established small-scale approaches. In particular, we sequenced a mixture of seeds pooled from several hundreds of *A. thaliana* genotypes from the 1001 Genomes Project (1001 Genomes Consortium, 2016), which can be used as a founder population for multiple evolution experiments (**Experiment 1**). We further constructed sequencing libraries of exactly two inbred genotypes using the Pool-Seq approach to assess the deviation in the allele frequencies from the expected 50% frequencies at positions where the genotypes differ (**Experiment 2**). We conducted varying poolings of genotypes and tissue types (i.e., leaf versus flower) to describe the effect of individual pooling and coverage in allele frequency inferences (**Experiment 3**). We ran a pilot “E&R common garden” experiment to test our methods in realistic outdoor settings and analyzed whether signals of rapid evolution could be detected in a few generations (**Experiment 4**).

### A Snakemake-based pipeline to streamline and parallelize frequency calling in Pool-Sequencing

To tackle the large amount of sequencing data that is needed to comprehensively test for rapid evolution across environments with Pool-Sequencing, we implemented `g`_r_`enepipe` (Czech and Exposito-Alonso, 2021), a pipeline based on the Snakemake workflow management system (Köster and Rahmann, 2012; Mölder et al., 2021), to process raw sequence data into variant calls and allele frequencies. We used `g`_r_`enepipe` to process the data from all our four experiments described below. Unless otherwise specified, we used `g`_r_`enepipe` v0.6.0, with the following tools in the pipeline: trimmomatic (Bolger et al., 2014) for read trimming, bwa mem (Li and Durbin, 2009) for mapping against the reference genome, and samtools (Li et al., 2009) for working with bam and pileup files. We furthermore employed several quality control tools that are built into `g`_r_`enepipe` to ensure that our sequence data is of sufficient quality (Andrews and Others, 2017; Ewels et al., 2016; Li et al., 2009; Okonechnikov et al., 2016). Note that `g`_r_`enepipe` furthermore offers variant calling, using tools such as BCFtools (Li, 2011), freebayes (Garrison and Marth, 2012), and the GATK HaplotypeCaller (McKenna et al., 2010). The exact tools and parameter settings used in each run of the pipeline are available at https://github.com/lczech/grenepilot-paper.

The `g`_r_`enepipe` automatization of single variant polymorphism (SNP) and their frequency calling allows us to test a number of variant filters and compare them in a standardized fashion. Specifically, we focused on quality controls related to:

1. Base quality filters based on Illumina PHRED scores.
2. Mapping quality filters to reduce the likelihood of false positive variant calls. These follow essentially the same curated filters of the PoolSNP pipeline used in the “Drosophila Evolution over Space and Time” resource (Kapun et al., 2021, 2020) .
3. Free discovery of genetic variants *vs* utilizing only 11,769,920 biallelic SNPs (out of 12,883,854) previously discovered in individual strains from the 1001 Genomes project (1001 Genomes Consortium, 2016) or a high-quality subset of the same genome set of 1,353,386 biallelic “bona fide” SNPs (Exposito-Alonso et al., 2019).
4. Coverage filters and minimal alternative allele counts to reduce sampling noise and sequencing errors (Kapun et al., 2021; Lynch et al., 2014).

The experiments described below make use of these filters, unless otherwise specified.

### A new efficient command line tool for population genetic statistics using Pool-Sequencing

To efficiently analyze Pool-Seq allele frequency data for thousands of population samples, we developed a C++ based command line tool called `g`_r_`enedalf` (Czech and Exposito-Alonso, 2022), which is able to parse .bam/.sam/.cram/.vcf/.pileup/.sync files, analyze allele counts and frequencies on the fly, and compute population genetic statistics implemented in the broadly used PoPoolation1 and PoPoolation2 (Kofler et al., 2011a, 2011b) along with new extensions of several unbiased statistics derived here and elsewhere (Hivert et al., 2018). The **Supplemental Mathematical Appendix** includes mathematical derivations and motivation of various unbiased corrections of Watterson’s *θ_W_* , Theta *π* , Tajima’s *D*, and *F_ST_* that account for two main sources of noise in Pool-Seq: the finite number of individuals pooled (*n*), and the finite coverage per base pair along the genome (*C*) (see cartoon **Fig. S1**). These are two nested Binomial samplings, where, for a polymorphic site in a population, we first have a chance of sampling *k* individuals carrying each occurring allele out of all *n* individuals pooled, which is proportional to the true allele frequency *f_A_* in the population. Then, after DNA sequencing, we have a chance of observing *c* reads in a pool of *C* (coverage) reads, which is proportional to the frequency of each allele in the pooled sample of individuals (*k/n)*.

The first parameter that we are interested is the genetic diversity, nucleotide diversity, or observed heterozygosity, for a given SNP in the genome, expressed as:

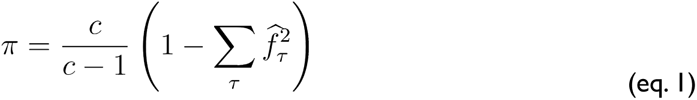

where 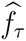 is the frequency representing each of the four possible nucleotide bases (*τ* ∊ ACTG), based on the nucleotide counts *C_τ_* of the reads in the pool, with total coverage *c = ΣC_τ_*. This is using Bessel’s correction for finite coverage; an additional correction of individual sample size n/(n-1) may also be applied, as described in the **Supplemental Mathematical Appendix**. Such a metric of diversity could be used to detect islands of low diversity appearing over time in E&R, which could be indicative of a selective sweep.

The second parameter of most interest is allele frequency differentiation between two spatial or temporal population samples, *F_ST_* , for which there are multiple definitions (**Supplemental Mathematical Appendix**). Following the same notation as the nucleotide diversity, Nei’s unbiased *F_ST_* for Pool-Seq can be defined as:

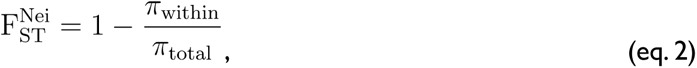

where the within, between, and total diversity can be calculated based on frequencies for two populations, coverages, and number of individuals pooled (indicated with subscripts *(1) (2)* for the two populations) as:

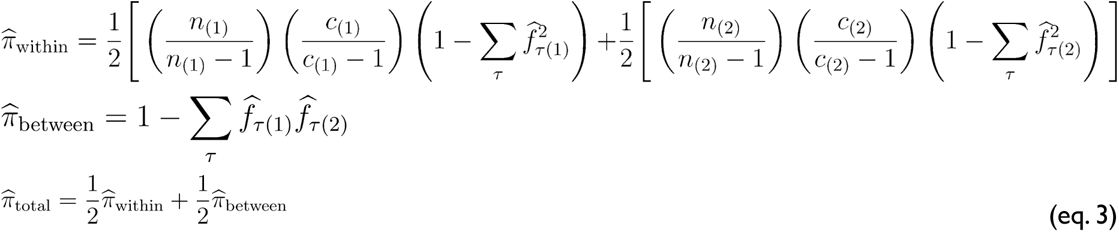

The above observed metrics are most useful for inferring processes within E&R experimental populations with known founders. When using Pool-Seq for natural populations, it may also be helpful to infer population parameters such as the population mutation rate *θ* (*4N_e_ μ*) from empirical diversity estimates such as *π* while accounting for Pool-Seq errors. The general strategy described in PoPoolation (Kofler et al., 2011a, 2011b) and reimplemented `g`_r_`enedalf` is described in detail in the **Supplemental Mathematical Appendix** .

### Experiment 1: Sequencing a seed mixture of 231 genotypes to characterize a diversity panel

#### Rationale

In this experiment, we established a genetically diverse panel of 231 *A. thaliana* natural accessions (i.e. seeds sourced from 231 different locales). We sequenced the mix of seeds (of roughly at equal proportions) of this panel to assess the ability of Pool-Seq to correctly recover genome-wide allele frequencies. This was the first step to use the seed pool for further E&R experiments (see below).

#### Setup

The founder seed mix for this experiment was sourced from the seeds of 231 genotypes, 229 of which are part of the 1001 Genomes Project (2016) and available from the Arabidopsis Biological Resource Center (ABRC) under accession CS78942 (https://abrc.osu.edu/stocks/465820), while the remaining 2 genotypes were sourced through the Israel Plant Gene Bank (https://igb.agri.gov.il/) under accession numbers 24208 and 22863 (**Dataset S1**). Seeds were pooled at roughly equal yet variable proportions based on weight (See **Dataset S1** for estimated numbers of seeds per ecotype). Note that despite differences in seed proportions in the seed mix, the intent of direct genome sequencing below is to capture these differences to establish an accurate allele frequency baseline.

#### Analysis

Eight tubes, each containing about 2,470 seeds (estimated based on weight) from the founder seed mix (**Table S1**) were homogenized using a FastPrep-24 (MP Biomedicals, Irvine, CA, USA). DNA extraction was done using a Qiagen DNeasy Plant Mini kit (Hilden, Germany) (**Supplemental Appendix I: DNA extraction**). One TruSeq library was prepared from each DNA extract. The eight TruSeq libraries were multiplexed and sequenced together on one lane of a HiSeq 3000 sequencer (Illumina, San Diego, California, USA). The total sequencing output was 9.54×10^10^ base pairs and the average genome-wide coverage was ∼500X across all seed pool sequencing data (**Fig. S5**). Raw sequence data were processed with our `g`_r_`enepipe` workflow (Czech and Exposito-Alonso, 2021) to trim and map the reads against the *A. thaliana* TAIR10 reference genome (Berardini et al., 2015; Lamesch et al., 2012). Subsequently, using our `g`_r_`enedalf` tool, we calculated the raw minor allele frequencies (MAF) at each biallelic position, based on bam/pileup files counting the ratio of reads containing either reference or alternative alleles (**Fig. S1**). Since users of Pool-Seq may utilize popular computationally efficient variant callers used in individual sequencing, we also ran `g`_r_`enepipe` with three different variant callers: BCFtools (Li, 2011), freebayes (Garrison and Marth, 2012), and the GATK HaplotypeCaller (McKenna et al., 2010). These tools are not primarily designed for calling variants (and their frequencies) from Pool-Seq data, but the resulting VCF file of each caller can be turned into a frequency table by extracting the Allelic Depth (“AD”) format field at each genome position for each sample, a process also implemented in `g`_r_`enedalf`. We also tried to run GATK HaplotypeCaller and freebayes using the average pool size as the ploidy options (`--ploidy 2470` and `--ploidy 2470 --pooled-discrete`, respectively, as well as `--pooled-continuous` in freebayes). Note that *A. thaliana* is diploid although inbred, but pooling ∼2,500 seeds would make the DNA library highly ploid from a computational point of view. These analyses resulted in prohibitively long runtimes even in cluster environments (GATK HaplotypeCaller) and large memory usage (freebayes), demonstrating these tools’ limited capabilities for analyzing large datasets and large pool sizes. We hence ran the three callers with default ploidy of 2 to study their artifacts in Pool-Seq applications, assuming that other researchers may be required to resort to these default settings.

#### Results

We conducted two comparisons, the first to quantify how well direct sequencing of pools of seeds captured the variation found in the 229 genotypes from the 1001 Genomes, and the second to study the technical artifacts generated by diploid SNP callers.

The first comparison is based on the raw frequency of alternative allele counts divided by coverage in the seed sequencing bam/pileup files against the same SNPs using the 229 columns in the 1001 Genomes VCF table corresponding to the genotypes mixed at roughly equal proportions in the seed mix (comparisons conducted with 1,353,386 *bona fide* SNPs with minimum alternative allele count >2). This yielded a high correlation and low deviation from the y=x correspondence line (**Fig. 1A**, Pearson’s *r* = 0.982*, SD* = 0.0214; for unfiltered comparison see **Fig. S3A-B**). Of note, comparing seed allele frequencies to all 1,135 individuals from the 1001 Genomes (i.e., not only the 229 included in the seed mix) shows a high density of alleles that are at low-to-intermediate frequencies in the 1001 Genomes but at low frequency in the seeds (**Fig. S3A,C**), likely indicative of a rare and highly divergent population group, the so-called relict accessions, comparatively underrepresented in our Pool-Seq subset of the larger set of 1001 Genomes (1001 Genomes Consortium, 2016). All in all, we were able to recover nearly all of the SNP variation present in the 229 individual founder genotypes with our Pool-Seq approach.

**Fig. 1.**
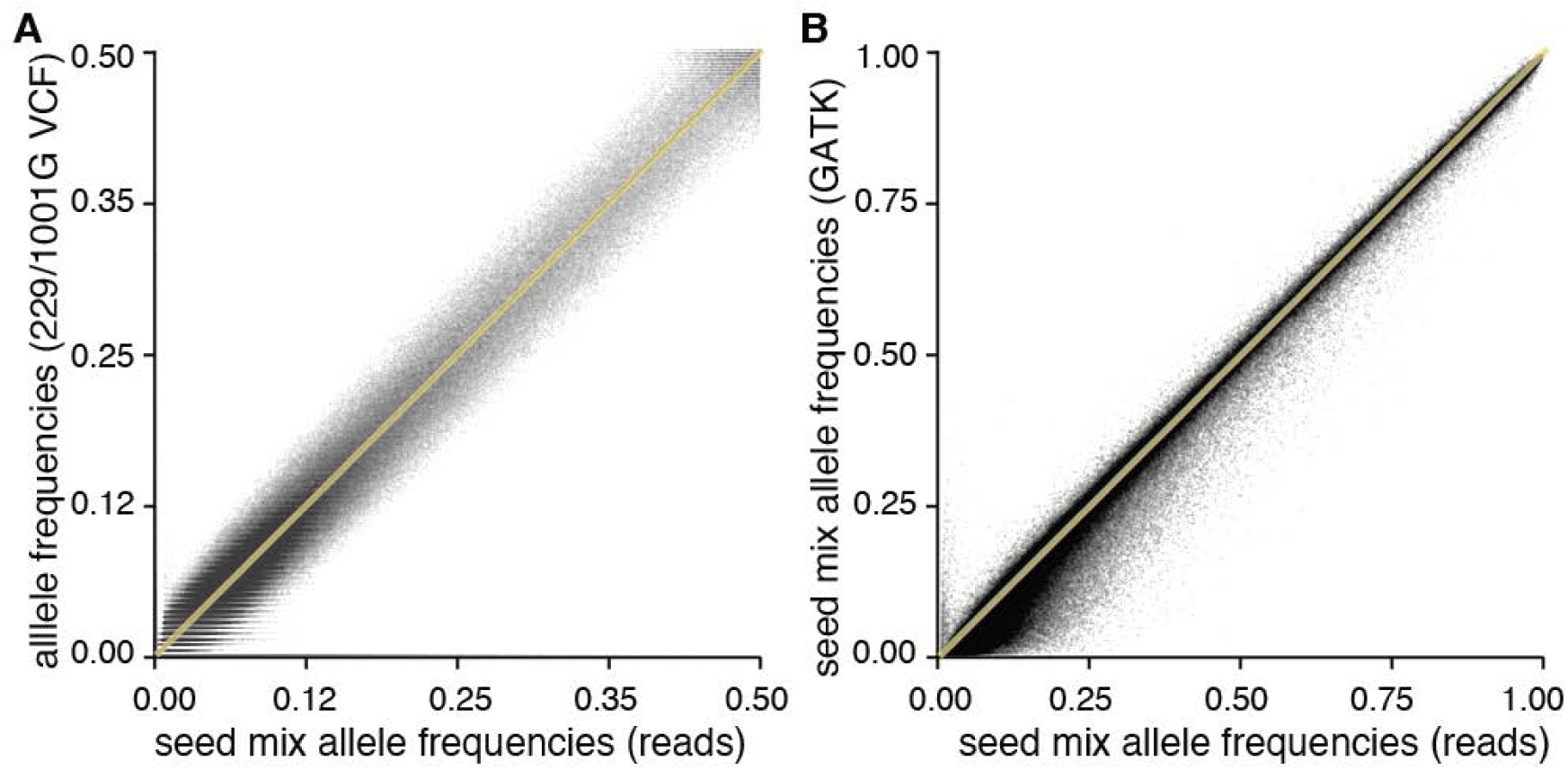
Direct sequencing of experimental founder seeds captures the 1001 Genomes variation. **(A)** Comparison of minimum allele frequencies directly estimated from ratios of bases in reads from sequencing the seed mix (x-axis) and allele frequencies calculated in silico from the 1001G VCF subsetted to the genotypes shared with the seed mix (y-axis) **(B)** Comparison of allele frequencies from the seed mix likewise directly calculated from ratios of bases in reads (x-axis) vs from the allelic depth (“AD”) VCF field after calling SNPs using GATK with default settings (y-axis). Yellow lines indicate y=x line.

The second comparison assessed the variation in allele frequency recovery from raw allele counts in bam/pileup files and standard diploid SNP callers. This revealed that certain tools generated frequency estimates upwardly or downwardly biased compared to the raw allele fraction from bam/pileup files (**Fig. S6-9**). This bias, especially for alleles found at low frequency (<20%), appears most dramatic in GATK HaplotypeCaller in its default diploid likelihood mode (**Fig. 1B, Fig. S6-9**). Further, very low frequency alleles (<4%) appear missing (**Fig. S10**). GATK, which is tuned for human SNP calling, aims to call genetic variants that fit the reference homozygote, heterozygote, or alternative homozygote scheme, and utilizes local genome realignment information to reject certain reads, which may be causing unexpected biases (see asymmetry in low-frequency SNPs in **Fig. 1B**, where only 1,353,386 *bona fide* SNPs with minimum count >2 were used). While correlations between raw allele frequency and SNP calling-based allele frequencies are typically high (*r* >0.99), deviations can be substantial without filters. For instance, the standard deviation of differences between GATK and raw allele ratio frequencies suggests deviations higher than 10% (SD_GATK_=0.094-0.204 depending on coverage cutoffs, **Fig. S7-9)**. BCFtools and freebayes appeared less biased and more consistent (SD_BFCtools_=0.051-0.156 and SD_freebayes_=0.043-0.097, **Fig. S6-9**). Based on these results, we recommend, despite their computational capacity and popularity, to avoid SNP callers designed for individual sequencing for Pool-Seq data. In conclusion, `g`_r_`enedalf` offers computational speed for computing diversity and differentiation statistics, generates frequency tables from raw sequencing reads, allows for data manipulations such as subsets or sample comparisons, and implements quality filters shown to provide appropriate frequency estimates, e.g., if evaluated in a set of *bona fide* SNPs (see Experiment 2 below) (Guirao-Rico and González, 2021).

### Experiment 2: Two-genotype analysis to understand biases of DNA contribution to pooled samples and sequencing noise

#### Rationale

One important assumption in population inferences based on Pool-Seq data is that each individual contributes an equal amount of sequencing reads. However, the deviation in DNA contribution by pooling organs from different individuals or entire individuals has not been tested in *A. thaliana* or other model plant systems (although it is common practice in *D. melanogaster* to directly pool whole flies, see for instance Tilk *et al*. (2019)). Instead, a typical approach in many state-of-the-art Pool-Seq experiments is to extract DNA separately from different individuals and subsequently pool equal amounts of DNA, an unfeasible approach when studying thousands or tens of thousands of individuals (Gautier et al., 2013; Rellstab et al., 2013; Roda et al., 2017). Whether flower organ sizes, such as those described in *A. thaliana* across ecotypes (Juenger et al., 2000), or cell ploidy differences via endoreplication in sepals (Robinson et al., 2018) have an effect in differential DNA contributions when pooling flowers, is unknown and could be manifested in deviations of allele frequencies. In Experiment 2, we sequenced a pool of two *A. thaliana* genotypes sampling one flower each (i.e., the smallest possible pool size *n*=2) and tested it against carefully quantified and pooled DNA isolates of the same two genotypes to assess the variation in DNA contribution.

#### Setup

To quantify the deviation in DNA content when pooling two flowers from distinct genotypes, we sequenced three replicates of two flowers each. The first genotype was the laboratory inbred strain Col-0, which was the type strain used to assemble the reference genome of *A. thaliana* (Lamesch et al., 2012). The second, a natural accession (inbred in greenhouse propagations) from the 1001 Genomes project (1001 Genomes Consortium, 2016), was RUM-20 (#9925), which differs from Col-0 by 1,007,560 SNPs according to the 1001 Genomes data (note that the average genotypic difference of any two genotypes is 400—600K SNPs; we hence picked a relatively divergent accession). These two ecotypes did not show visible flower size differences, but were not chosen based on their flower size differences. To compare the pooled flower method with the conventional method where DNA is pooled at equal proportions, we extracted DNA from a leaf of a Col-0 individual and a leaf of a RUM-20 individual, and generated three DNA replicates via equal pooling by DNA concentration before library preparation (**Fig. 2A**, **Table S2**). DNA was extracted with the CTAB method and processed into whole-genome sequencing libraries using a modified Nextera protocol (**Supplemental Appendix I: DNA extraction** and **Library preparation**).

**Fig. 2.**
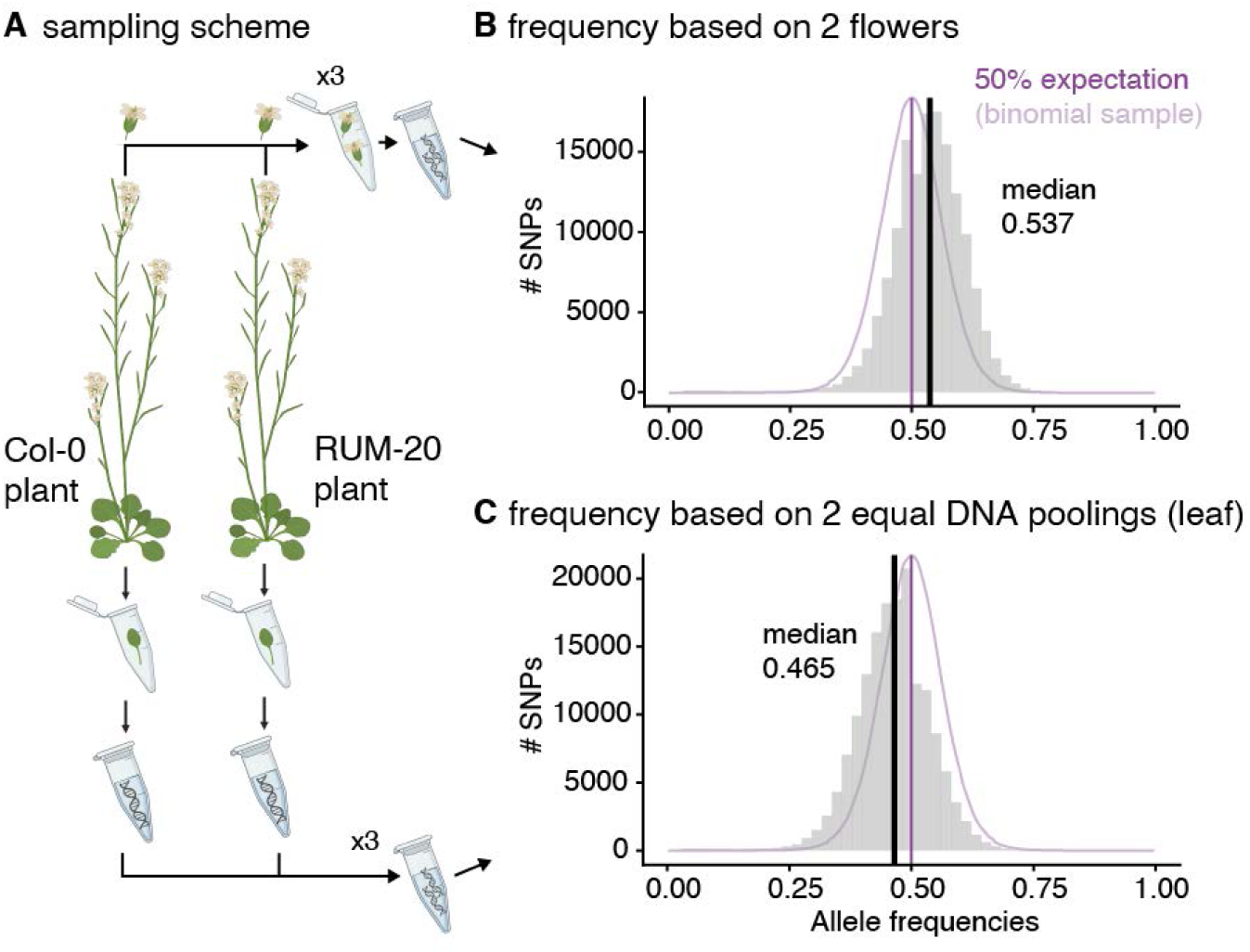
Experimental design (Exp. 2) to test the relative contribution to DNA sequencing output. **(A)** Flower and leaf tissues were sampled from two genotypes, Col-0 and RUM-20. Three replicates of two flowers were collected (the Pool-Seq method) while leaves were collected individually (conventional method). Leaf DNA was pooled at equal quantity to create three replicates of DNA input for library preparation. **(B)** Distribution of allele frequencies in one of the three replicates of the directly extracted and whole-genome sequenced 2-flower pools (see all replicates in **Fig. S14**; allele frequencies of SNPs that passed mapping quality filters, had a minimum minor allele count >2, and were present in the 1,353,386 *bona fide* SNPs). **(C)** The equivalent of (B) for two separate leaf DNA extracts carefully pooled at equal concentration.

#### Analysis

Assuming that both pooled individuals from the two inbred lines are indeed homozygous, one would expect polymorphic alleles to be at exactly 50% proportion if the tissues of both individuals contributed exactly equal amounts of DNA. Mean deviations from 50% would indicate differences in DNA content and/or mapping bias if deviations are systematic for one genotype. In addition, variations in the extent of deviation across replicates would indicate sampling noise due to limited DNA sequencing coverage. To test this, we again trimmed and mapped genome-wide reads with our `g`_r_`enepipe` workflow, and computed frequencies from bam/pileup files with `g`_r_`enedalf`, as described in Experiment 1.

#### Results

This proof-of-concept analysis provided a number of clues on the power and potential biases of Pool-Seq to study population evolution in real time. Firstly, as the expected frequencies of polymorphic sites are around 50%, we were able to detect many low frequency alleles that are most likely artifacts (**Fig. 2B-C**). This could not be done while sequencing large populations of seeds or flowers because we expect many true low frequency alleles. We show that a lack of filters for coverage, mapping quality, and most importantly, minimum alternative allele count, leads to a majority of calls of polymorphic sites likely representing artifacts (e.g., 90% of unfiltered SNPs may be false positives, see **Fig. S11A**; these filters were applied in all experiments). We therefore implemented stringent filters for mapping quality (samtools option ‘`-q 60`’), base quality (option ‘`-Q 30`’), and matching of forward/reverse read mapping (options `‘--rf 0x002 --ff 0x004 --ff 0x008’`), and minimum allele counts in the bam/pileup file of reads (MAC>2). In combination with a filter for the *bona fide* 1,353,386 SNPs, these filters led to the expected distribution of allele frequencies around 50% with some of the remaining variation likely explained by the binomial sampling variance caused by limited coverage (**Fig. 2B-C, Fig. S11**). Third, we were able to show that the deviation of average allele frequencies from 50% was small across the three flower pool replicates (2.2% deviation in frequency from 50%, see **Fig. S13D-F**), and of similar magnitude as the deviation measured in the DNA pools generated from three DNA isolates carefully pooled with equal concentration after DNA normalization (1.5%, deviation in frequency from 50%, see **Fig. S13A-C**). This suggests that uncontrolled factors (such as flower size, endoreplication and ploidy, differential tissue grinding) minimally affect DNA contributions of flowers, and that their magnitude is comparable to variable DNA contributions from individual samples even after DNA normalization. Such small deviations become statistically diluted when pooling large numbers of individuals (Lynch et al., 2014). For instance, for 100 flowers, errors would range from 0.0004 to 0.1% for allele frequencies from 1% to 50%. This is in agreement with previous Pool-Seq experiments with whole *D. melanogaster* flies, which indicates that allele frequency estimation per population requires 100 individuals at 50X coverage for virtually-perfect allele frequency retrieval (Gautier et al., 2013). In summary, the Pool-Seq approach using large numbers of *A. thaliana* flowers, sampling one flower per individual, should provide highly reliable allele frequency inferences in E&R experiments.

### Experiment 3: Combinatorial experiments of pool sizes and tissue type sequencing to determine optimal sampling schemes

#### Rationale

In this experiment, we evaluated the ability of Pool-Seq to recover correct allele frequencies from pooled samples made up of 5 to 100 flowers (one per individual) and leaves sampled from *A. thaliana* plants. We studied whether (A) individual leaf DNA extraction and library preparation with equal DNA input and (B) pooled flower DNA extraction and library preparation without DNA normalization produce comparable population estimates.

#### Setup

We grew a mixture of seeds of 231 genotypes mixed roughly at equal proportions (**Dataset S1**) in 2,500 pots with one individual each (replicating similar conditions of large evolving populations outdoors, see Experiment 4). We then selected 50 random plants for our test. Flowers were sampled from different subsets of these 50 plants to assess the effect of increasing the number of individuals randomly sampled: 5, 10, 25, 50, and 100 flowers (**Fig. 3**; for 100 flowers, we included 2 flowers from each of the same 50 plants). For the same plants for which flowers were collected, we also removed, and separately stored, one leaf per plant for independent DNA extraction.

**Fig. 3.**
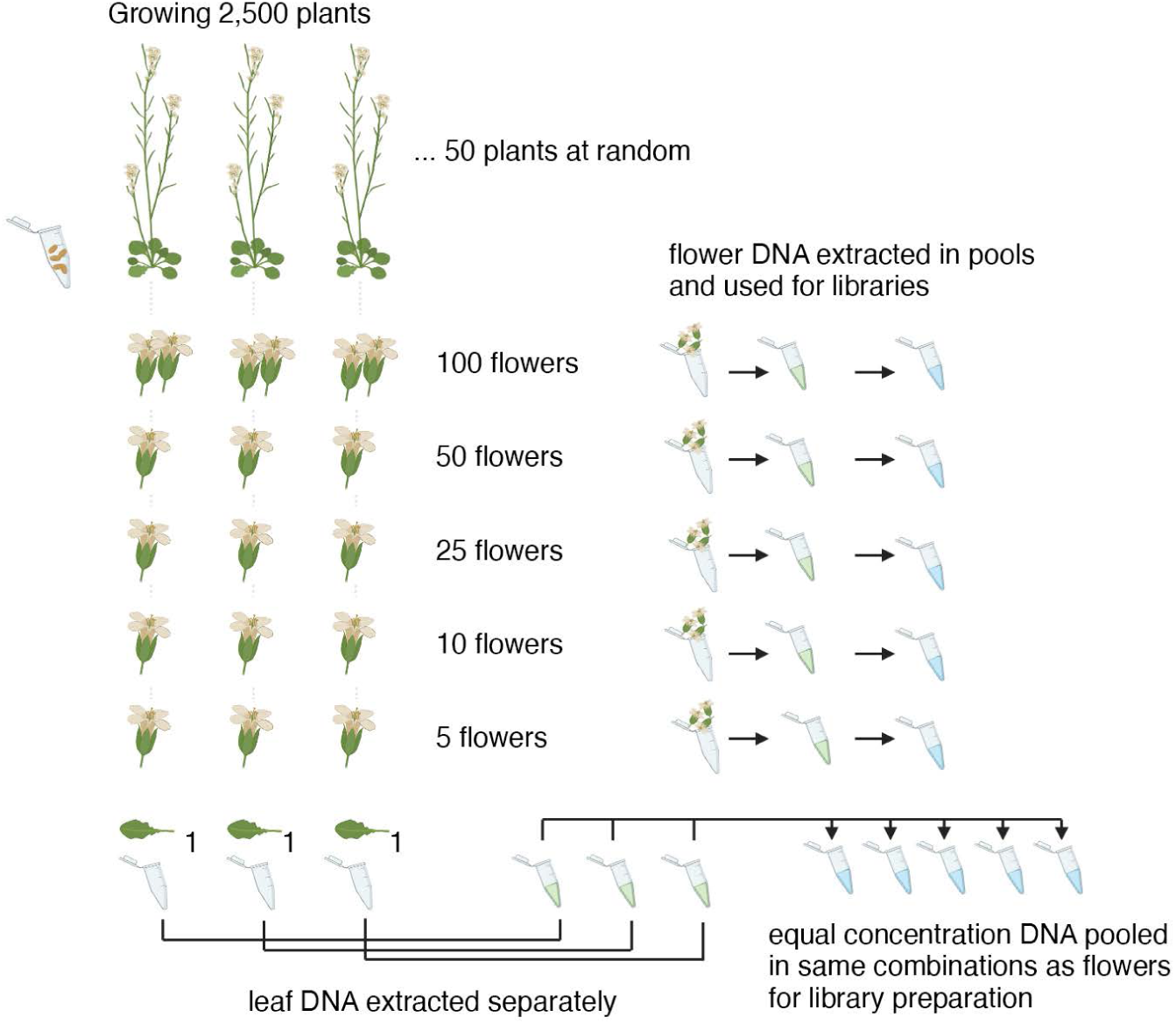
Experimental design (Exp. 3) to test Pool-Seq with plant flowers. A total of 50 plants sown at random from 231 diverse genotypes of *A. thaliana* (**Table S2**) were individually grown and sampled in different combinations and in replication (5, 10, 25, 50. In the 100 flower sampling, 50 individuals sampled twice) (**Table S4**).

Tissue grinding, DNA extraction and library preparation steps are described in **Supplemental Appendix I: DNA extraction and library preparation**. Leaf DNA pooling was done for the same individual combinations for which flower subsamples of 5, 10, 25, and 50 individuals were taken and pooled (see **Table S3-4** and **Fig. S2** for combinations). Therefore, we expect the allele frequencies of the equimolar pool of leaf DNA and that of the flower extracts to be close to identical (as in Experiment 2), unless scaling the Pool-Seq method to many individuals incurs systematic biases.

#### Results

We calculated the correlation between allele frequencies recovered from pools of flowers and equal leaf DNA pools both originating from the same sets of plants. Because noise decreases with both increasing numbers of individuals and increasing sequencing coverage, we leveraged the variation in coverage along the genome to compute correlations in increasing coverage bins. Frequencies were highly correlated (*r* > 0.98) for all combinations as long as coverage was over 50X (**Fig. 4**). The small mean relative frequency differences (<15%) of alleles at medium (∼40X) coverage for virtually all pairs of flowers or DNA pool libraries suggests that, even when there are small experimental pooling errors (**Fig. 2B-C**), the large number of sequenced individuals dilutes errors (**Fig. 4A** ) (Lynch et al., 2014).

**Fig. 4.**
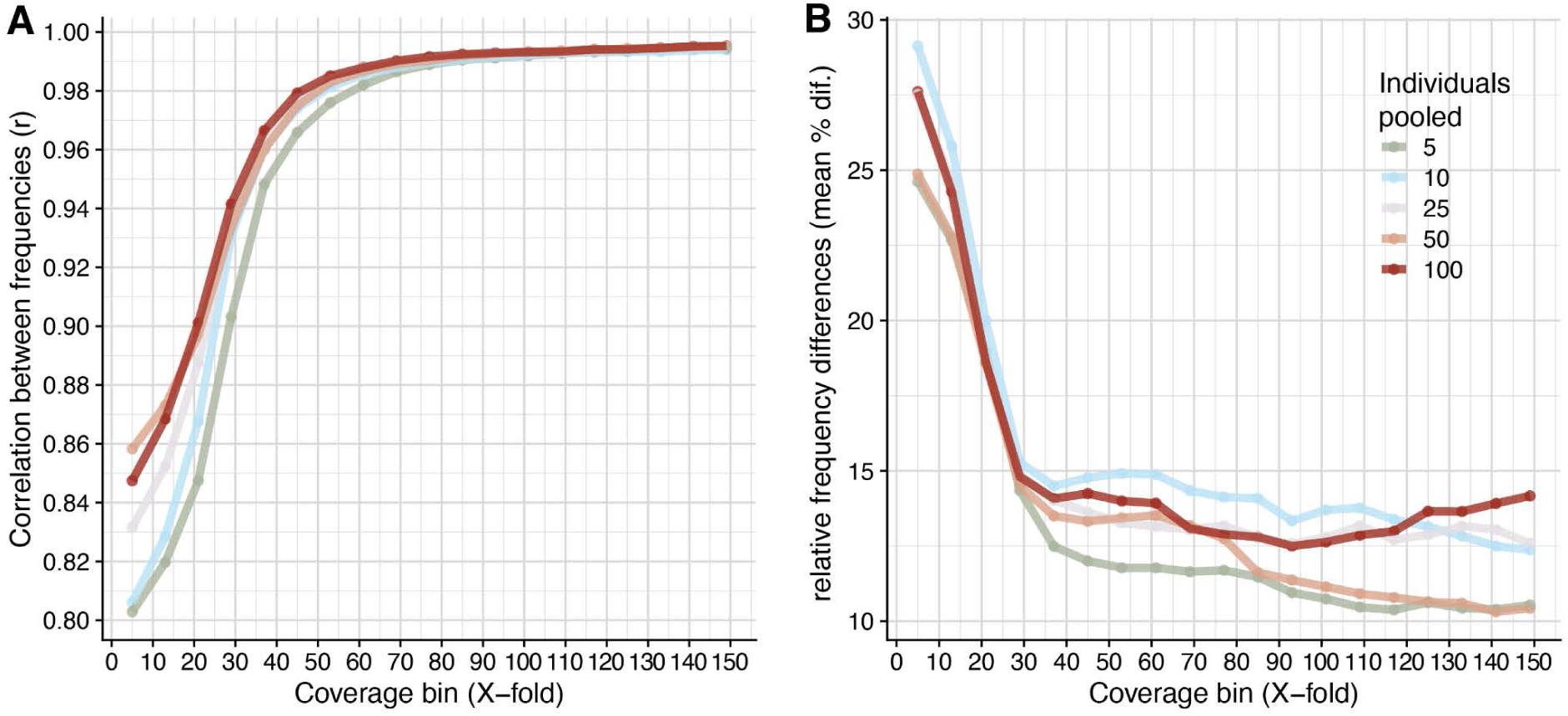
Correlation between allele frequencies estimated from direct sequencing of pooled flowers *vs* individual DNA extracts pooled at equal concentration. Genome-wide allele frequency comparisons between the same set of 5,10, 25, or 50 individuals as estimated from directly extracting DNA and sequencing from pooled flowers and from sequencing of pooled independent DNA extracts. (**A**) Pearson’s correlation between allele frequencies across coverage bins (all alleles with minimum alternative allele count >2 and represented in the *bona fide* 11,769,920 set (further subsets based on other quality thresholds did not provide enough data points for coverage breakdown). (**B**) Relative % error of the difference between flower pools and DNA pools across coverage bins.

### Experiment 4: Multi-year field experiment to showcase the power of Pool-Seq to track rapid evolution

#### Rationale

Ultimately, the cost-effective and scalable Pool-Seq approach is designed to track evolution of populations through time. To showcase its strengths, we conducted an outdoor experiment over two growing seasons starting from a large population of diverse *A. thaliana* genotypes.

#### Setup

This experiment was performed in an experimental field at the Max Planck Institute of Biology campus (<u>48.537723, 9.058746</u>, Tübingen, Germany, **Fig. 5**-**6**), using a seed mix of 451 natural genotypes generated from 2 plants of each genotype and 10 siliques each (ca 90,000 seeds). This set largely overlaps with the 1001 Genomes genotypes, and therefore the starting allele frequencies are known (available in ABRC stocks under accession CS78942, https://abrc.osu.edu/stocks/465820, **Dataset S1**). The seed mixture was split in nine tubes which were sown at three different time points (November 2014, February 2015, and March 2015) in three independent 1×1 m^2^ plots (**Fig. 6A**). This design was used to reduce the chance of disturbance events occurrences that would impact germination. After the first generation had dispersed seed during late spring in the field and synchronized with the local climate and photoperiod, soil samples were collected prior to the second season’s natural germination (early fall 2015) and transferred to an indoor greenhouse with optimal conditions for the species, to enable germination, survival, and reproductive success for as wide a set of genotypes as possible. From the three plots, 56, 69, and 101 adult plants were sampled from the growth chamber as a baseline (Time 0 in **Table S6 and** **Fig. 5**). From the field plots, in the following spring, 164, 415, and 593 surviving and reproducing adults of the second generation were sampled for sequencing at 3 different time points to capture the entire temporal window of flowering (see number of adults per time sample in **Table S6 and** **Fig. 5**). We aimed to sample one flower per individual, paying attention to sample from small to large plants uniformly. The flowers collected at Time 1, 2, and 3 were sequenced separately. A total of 1,398 individuals were sequenced in 12 pools of replicate x time point combinations (**Table S6**). We used the unbiased pool-sequencing estimator for *F_ST_* (Nei), as described in the **Supplemental Mathematical Appendix** and as implemented in `g`_r_`enedalf`, to determine genome-wide patterns of *F_ST_* across all combinations of replicates and time points accounting for pool size (**Table S6**) and genome-wide variation in coverage.

**Fig. 5.**
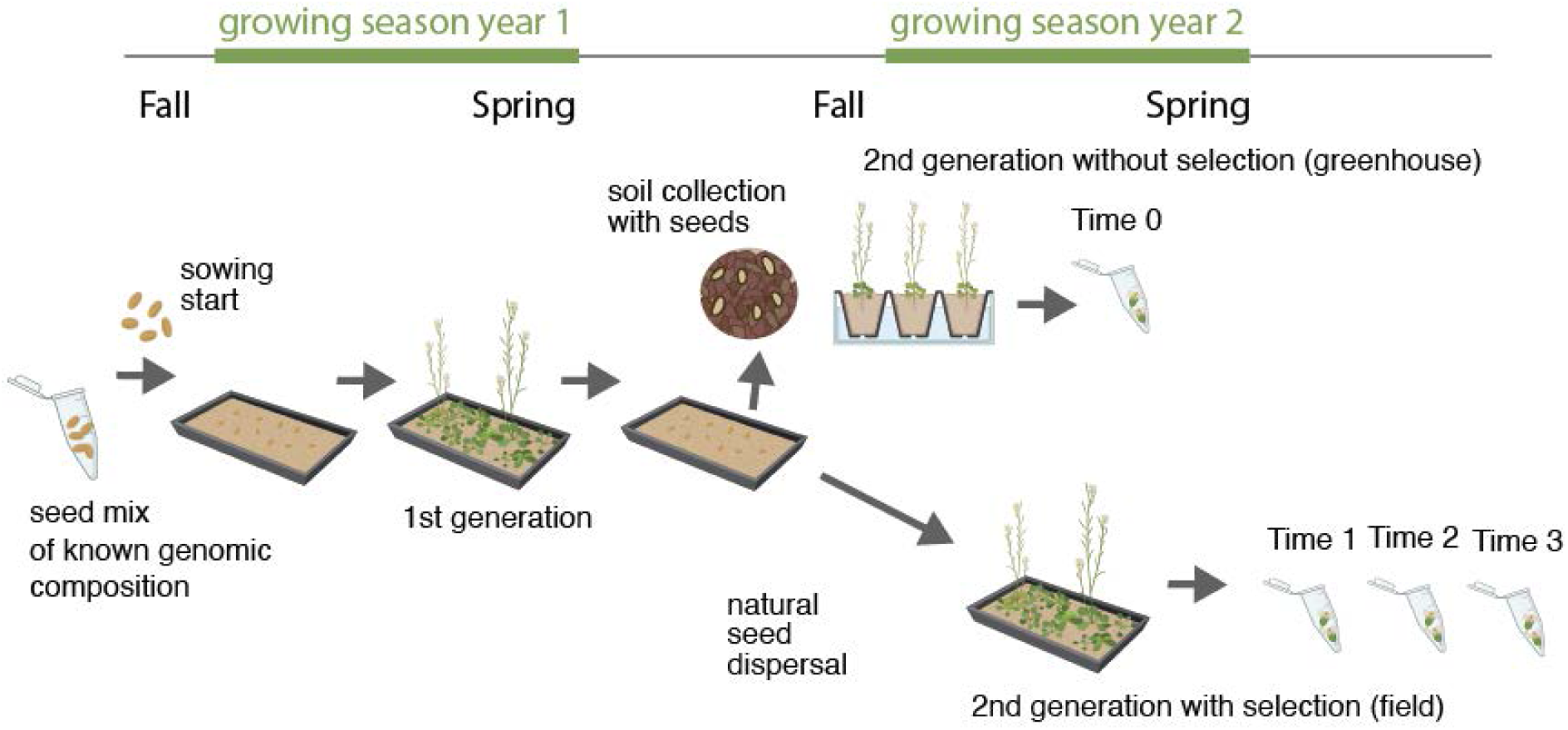
Design of the field experiment (Exp. 4) A plot was set up containing a mixture of seeds of 451 natural genotypes mixed at equal proportions. After an entire generation of growth in the field with natural seed dispersal, two parallel samplings were conducted. One sampling of soil was conducted in fall prior to natural germination and then planted in a greenhouse and subject to environmental conditions favorable for germination and growth to limit natural selection. The second sampling was conducted in spring after natural germination had occurred and plants were exposed to natural selection that could have led to mortality and survival of different genotypes. Experimental populations outdoors were sampled three times due to longer flowering periods in outdoor conditions.The whole experiment was replicated three times in parallel.

**Fig. 6.**
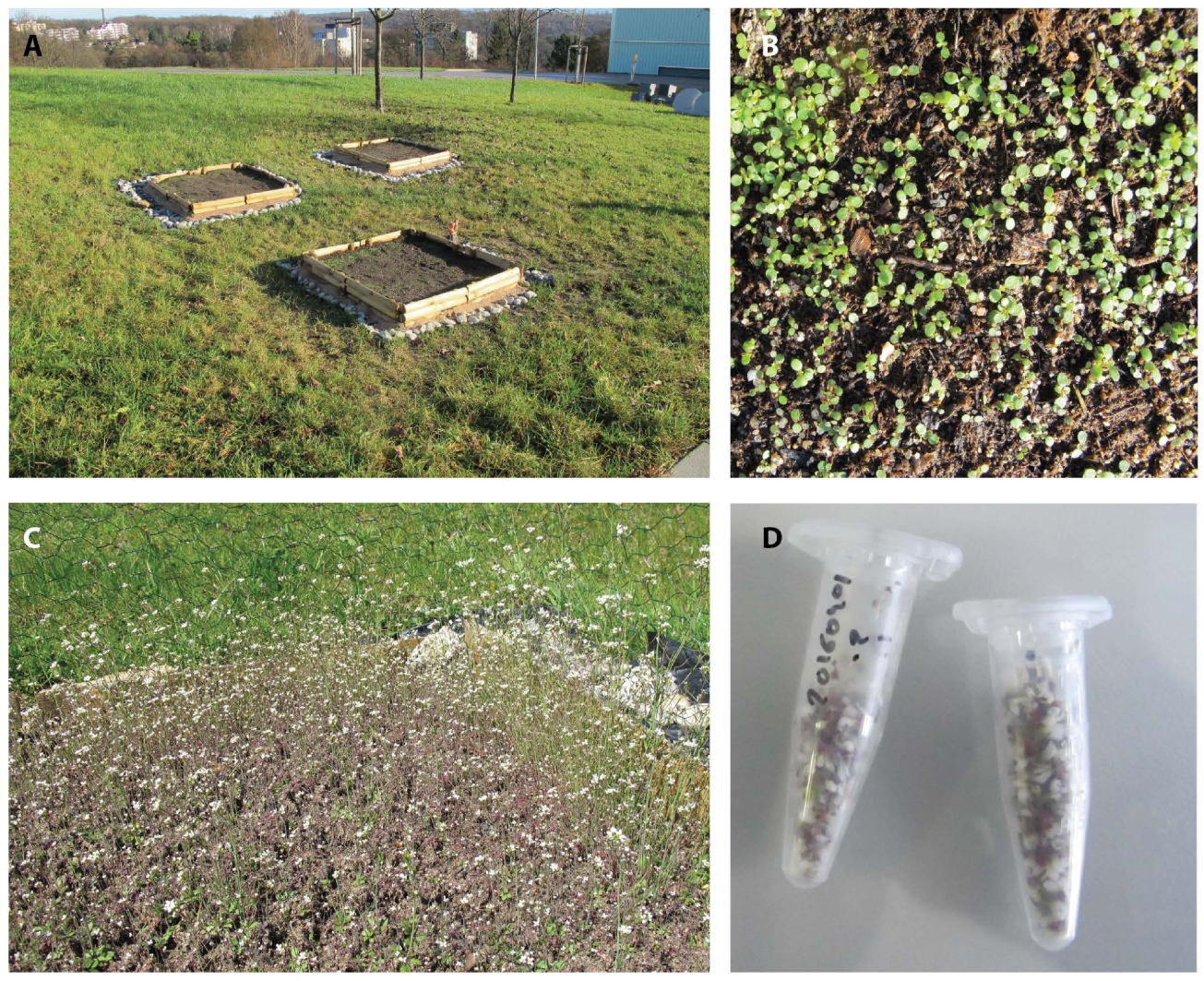
Photos of the field experiment (Exp. 4) (**A**) Setup of 3 population replicates. (**B**) Close-up of germinating seedlings. (**C**) Abundant flowering shown in one of the replicate plots. ( **D**) Sampled flowers for Pool-Sequencing.

#### Results

Plants successfully established in dense patches in the experiment (**Fig. 6B-C**). Tens of thousands of seedlings were observed per plot replicate, which theoretically should enable efficient natural selection (Charlesworth and Charlesworth, 2010). We observed genetic differentiation based on *F_ST_* between the baseline (*Time 0*, offspring of the first generation read in the greenhouse) and flowers of surviving individuals of generation 2 (*Time 1,2,3*) was higher (*F_ST_* ≈ 0.0023, **Fig. S16**) than differentiation between several independent DNA extractions of subsets of the founder seed mixes (Exp. 1, *F_ST_* ≈ 0.0006, **Fig. S14**). Although we think such genome-wide patterns are likely mostly driven by drift in the wild, a scan along the genome identified several *F_ST_* peaks between *Time 0* and *Time 1-3*, revealing genomic regions that diverged above the background noise level (**Fig. 7, Fig. S17,19**).

**Fig. 7.**
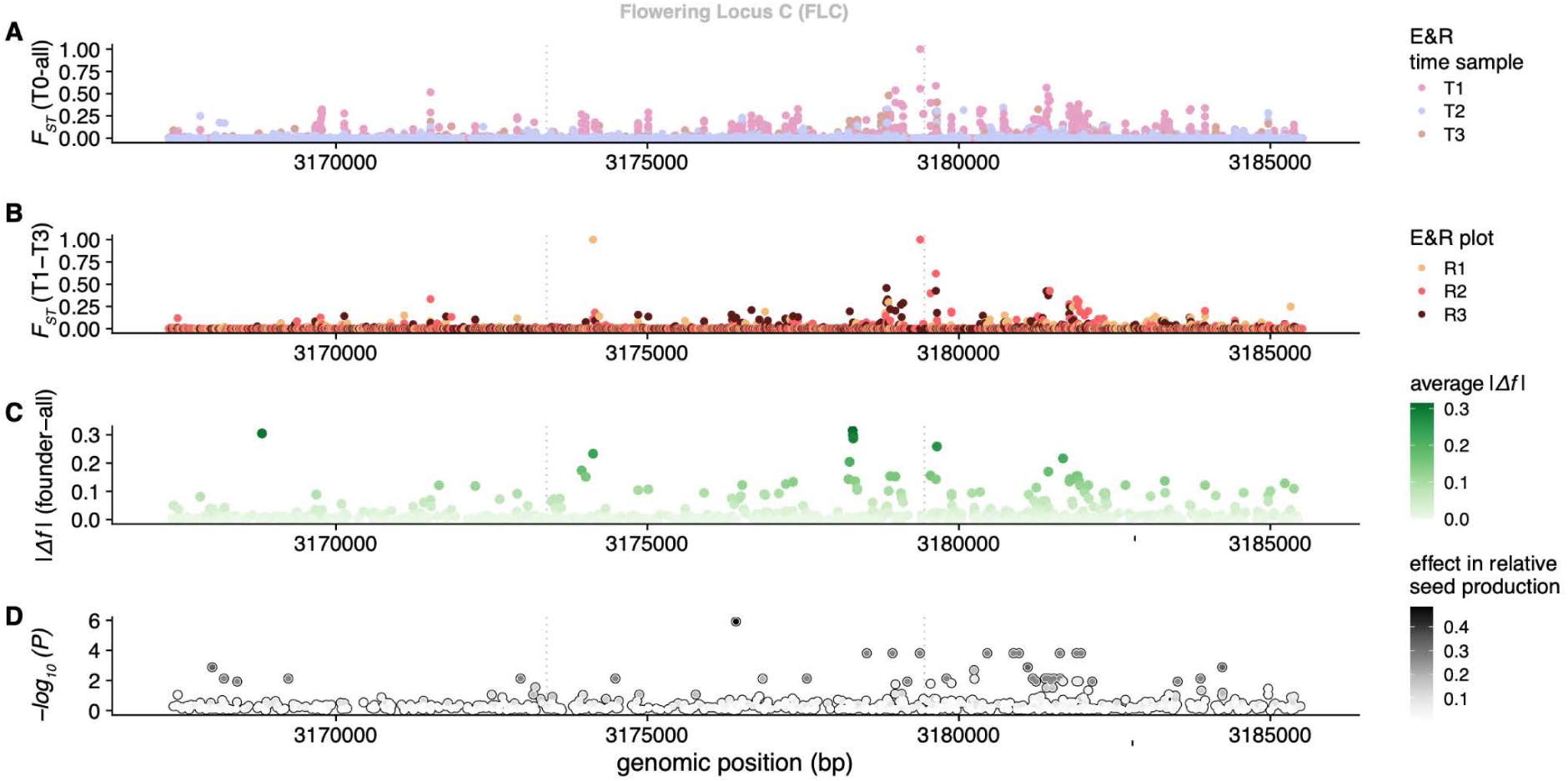
Temporal allele frequency change in a multi-year Evolve & Resequence experiment compared to fitness effects in a common garden experiment in the FLC region. (**A-B**) Temporal allele frequency differentiation (F_ST_, using our unbiased pool-seq Nei estimator) in the Flowering Locus C region on chromosome 5 showing peaks of differentiation around the first exon and the upstream promoter region of the gene (positions around 3,180,000; note the protein coding strand is the reverse strand). (**A**) Differentiation between the baseline “without selection” (Time 0) and the flower samples of surviving adults in nature at three time points (Time 1-3). (**B**) Differentiation between the earliest and latest flowering cohort for the three replicate plots. (**C**) Average allele frequency change between the founder seed mix (one generation prior to Time 0) to adults sampled in generation 2 (Time 1-3). (**D**) Genome-Wide Association between genetic variants in the 1001 Genomes and outdoor seed production in a common garden experiment (Exposito-Alonso et al., 2019) 1.51 kilometers away from the Evolve & Resequence experiment in (A-B).

One of the observed peaks is localized in chromosome 5 near the gene *FLOWERING LOCUS C* (*FLC*, AT5G10140), encoding a MADS-box transcription factor and master regulator of flowering time. The region with elevated *F_ST_* is located 5’ of the transcription start site of *FLC* (ca. -2.5—0.5K, **Fig. 7**), suggesting that variation in the promoter region was under some form of natural selection in these experiments and thus shifted in allele frequency. Average per-SNP *F_ST_* from *Time 0* to all other time points was higher within the approximate promoter region compared to the rest of the genome (mean [95% quantile] = 0.002964 [0.041188] in promoter vs. 0.009110 [0.010020] outside combining all three replicates; Wilcoxon signed-rank test *P<* 2.2 × 10^-16^, t-test *P=* 2.8 × 10^-11^) (**Fig. 7A**). That the same *F_ST_* peak is recovered by comparing two cohorts in the flowering seasons, *Time 1* vs. *Time 3* , further suggests variation in this genomic region may play a role in determining early vs. late flowering (**Fig. 7B**). Not only that, but raw allele frequency changes from the starting mix of 451 natural genotypes to all sampled flowers (1,172) two generations after the start of the experiment (**Fig. 7C**).

We leveraged the fact that Experiment 4 was conducted in parallel to a previous common garden experiment 1.51 km away (48.545809, 9.042449) with similarly rich and highly-overlapping *A. thaliana* genotype sets (Exposito-Alonso et al., 2019). In the common garden, each genotype was (individually) scored for an estimated number of seeds per plant that reached adult reproductive stage. Using an imputed matrix of the 1001 Genomes (http://arapheno.1001genomes.org, https://aragwas.1001genomes.org) and a Linear Mixed Model (Kang et al., 2008), we conducted Genome-Wide Associations (GWA) to identify genetic variants that explained variation in seed set per plant (**Fig. 7D**), specifically in the “thp” condition of that experiment: Tübingen, high rainfall, population replicate, (Exposito-Alonso et al., 2019) (For similar evidence in the “mli”: Madrid, low rainfall, individual replicate, see **Fig. S19**). This common-garden-scored fitness and GWA approach is one of the most direct ways to quantify natural selection driven by a specific environment (Exposito-Alonso et al., 2019; Gompert et al., 2017). It is expected that genetically-based fitness differences among plants would lead to genotype and allele frequencies changes over time (although such multi-generational experiments are not often conducted). As expected, we also found an overlap between the above peak of temporal *F_ST_* allele frequency differentiation in Experiment’s 4 E&R and moderate fitness-associated SNPs in the parallel common garden (**Fig. 7D**), with an average of fitness effect sizes significantly elevated within the same region observed above (Wilcoxon test *P =* 0.0313). The fact that flowering time, manually scored in the parallel common garden, was negatively correlated at the plant level with relative seed production (Spearman’s rank correlation *r* = -0.404, S = 31048965, *P* < 2.2 × 10^-16^) and survival (*r* = -0.187, S = 26399658, *P* = 2.074 × 10^-5^) further supports our finding that natural selection may have driven frequency changes in alleles in the *FLC* locus in our multi-year E&R field experiment. While the signal in the *FLC* locus is more readily interpretable, and is thus a helpful example to illustrate the application of our methods, this region is far from being the only region displaying strong temporal differentiation (**Fig. S18**). Multiple regions had *F_ST_* > 0.2 and showed parallel patterns in three or more replicates or temporal samples of flowers (**Dataset S2**). Although some genes involved in disease or dehydration responses are suggestive, most difficult-to-interpret peaks will deserve more attention in future studies. All in all, our experiment fulfills the purpose of testing the ability of a simple and cost-effective Pool-Seq approach to detect rapid evolution of plants subject to strong natural selection pressures at resolutions comparable even to those of time-intensive and costly common garden experiments and Genome-Wide Association studies.

### Discussion and Outlook

The paradigm that evolution is a slow process is being challenged by more and more evidence from experiments with both animals and plants that allele frequencies within populations fluctuate or change in the span of seasons or decades following environmental changes (Bergland et al., 2014; Franks and Weis, 2008). Scalable whole-genome sequencing approaches based on Pool-Seq (Schlötterer et al., 2014) have enabled the generation of population genomic datasets across continental scales such as “*Drosophila* Evolution over Space and Time” (DEST) (Kapun et al., 2021) (https://dest.bio) or large-scale multi-generational Evolve & Resequence experiments with *D. melanogaster* (Rudman et al., 2021). Projects of a similar scale for plants are currently rare, a notable exception being the barley composite cross long-term evolution experiment initially developed by Harlan and then continued by Jain and Allard (Allard and Jain, 1962; Suneson, 1956). Here we present Pool-Seq laboratory protocols and new efficient software implementations which are scalable to high-throughput, longitudinal experimental evolution studies of thousands of plant populations at low cost.

We here presented an open-source and streamlined frequency calling pipeline that automatically downloads, checks, and runs all the required software tools from raw fastq file to a frequency table of Pool-Seq samples (Czech and Exposito-Alonso, 2021). Such a reproducible pipeline also facilitates parallel runs with different pipeline parameters for tool benchmarking and quality controls for tool parameter comparisons. We show that if a set of *bona fide* SNPs is already known for the species, as is the case with *Arabidopsis thaliana’*s 1001 Genomes Project catalog (1001 Genomes Consortium, 2016), estimation of allele frequencies from mapped reads is successful without the need for sophisticated SNP callers to identify new variation, as long as there is sufficient coverage and quality filters are implemented (Guirao-Rico and González, 2021; Tilk et al., 2019). Two cases may benefit from further tool implementation in `g`_r_`enepipe:` In the absence of *bona fide* SNPs, Pool-Seq-specific likelihood or Bayesian SNP callers such as SNAPE are ideal to discover new SNPs while reducing false positives (Guirao-Rico and González, 2021). In the presence of ultra-low coverage sequencing, if individual sequencing of founders is available, allele frequency estimates can be further improved using simulations and linkage disequilibrium information based on the tools HARP and HAFpipe (Kessner et al., 2013;Tilk et al., 2019).

To enable faster and more user-friendly Pool-Seq-based evolutionary analyses at scale, we have developed `g`_r_`enedalf`. This tool re-implements the now-classic PoPoolation1/2 software (Kofler et al., 2011a) in C++ from the ground up, expands its functionality and types of compatible input file formats, and adds unbiased F_ST_ estimators for pool sequencing data, as described in the **Supplemental Mathematical Appendix**. Speed improvements in the order of ∼100X, in combination with multi-threaded scaling for powerful computers, now enable conducting, for instance, pairwise *F_ST_* calculations among thousands of samples in hours rather than months (Czech and Exposito-Alonso, 2022).

With these bioinformatic improvements in hand, we show that direct whole-genome sequencing of a mixture of seeds can properly characterize the standing genetic variation of a hypothetical starting pool of founder individuals for an E&R experiment. Further, direct sampling of flower tissues (or similarly-sized organs or leaf punches) also enables efficient genetic tracking of plant populations with tens of thousands of individuals over time—a scale currently not feasible for experiments with separate individual DNA extracts or library preparations (Fracassetti et al., 2015; Gautier et al., 2013; Rellstab et al., 2013; Roda et al., 2017). This sampling method potentially provides an alternative experimental design to common garden experiments, and its simplicity would potentially facilitate citizen-science real-time evolution projects in large organisms.

Finally we showcase that the described Pool-Seq protocols can be applied in large outdoor E&R experiments using *A. thaliana* seed resources. The fact that linkage decays surprisingly fast in *A. thaliana* (**Fig. S4**) (Kim et al., 2007)—probably owing to a ∼2-16% outcrossing rate that shuffles enough standing genetic variation (Bomblies et al., 2010; Platt et al., 2010)—may enable identification of narrow mapping regions containing adaptive loci using E&R, perhaps even narrower than what Genome-Wide Associations can currently achieve (**Fig. 7**) (1001 Genomes Consortium, 2016; Atwell et al., 2010). The success of Experiment 4 motivates the use of this approach at a larger scale, and seems to provide a genomic sensitivity similar to labor-intensive common garden experiments that are confined to a few environment (Agren and Schemske, 2012; Exposito-Alonso et al., 2019, 2018; Fournier-Level et al., 2011; Manzano-Piedras et al., 2014) ,

Despite the intriguing and complementary association between fitness effect sizes in common garden experiments and allele frequency changes in our E&R (Experiment 4), the rapid evolutionary signals inferred here are limited to the single environment studied. To comprehensively study rapid evolutionary adaptation across climates using E&R, we have initiated a project called “Genomics of rapid Evolution to Novel Environments” network (GrENE-net), which is a large-scale extension of Experiment 4 presented here. This internationally distributed E&R GrENE-net project involves 45 field sites (https://grenenet.org), was started from the same seed mix of Experiment 1, and has been conducted from 2017 until 2022 (the time of writing)—featuring the largest temporal and spatial scale among known Evolve & Resequence experiments. The accumulating sequencing data, expected to exceed 5Tb and over 2,500 population samples, should enable better temporal and spatial tracking of rapid evolution and understanding of climate × genotype × fitness interactions than any previous large-scale common garden experiment (Exposito-Alonso et al., 2019; Fournier-Level et al., 2011; Lovell et al., 2021). It is our hope that the GrENE-net experiment will enable researchers to establish a direct link between environment and natural selection at the allele frequency level, stimulate theoretical development in evolutionary genetics, and empower plant biologists’ search for the genetic basis of adaptation. If biologists wish to forecast plant responses under changing climate conditions, long-term and highly spatially replicated E&Re datasets such as this one will be paramount.

## Additional Information

### Data and Code availability

Reads were deposited at NCBI SRA with accession number: <TBD upon publication>. Genomes of founder populations are available as part of the 1001 Genomes project: http://1001genomes.org/data/GMI-MPI/releases/v3.1/. Scripts of the analyses in this manuscript are available at https://github.com/lczech/grenepilot-paper, which contains all settings used for the runs of our `g`_r_`enepipe` (Czech and Exposito-Alonso, 2021) workflow for variant calling, as well as all python and R scripts for the figures presented here (`g`_r_`enepipe` is available at https://github.com/moiexpositoalonsolab/grenepipe). Genome frequency manipulations and Pool-Seq-corrected population genetic statistics are implemented in `g`_r_`enedalf` (Czech and Exposito-Alonso, 2022) (available at https://github.com/lczech/grenedalf), and detailed equations and differences among estimators are described at https://github.com/lczech/pool-seq-pop-gen-stats.

## Acknowledgements

We thank Robert Kofler and Nicola De Maio for discussing their PoPoolation and PoPoolation2 implementation details. We further thank the members of the Moi Lab, Mark Bitter, and Dmitri Petrov, for feedback on the manuscript and analyses.

## Author contribution

MEA, FV, JFS, conceived the project after initial discussions with DW. MEA and DW acquired financial support for the project leading to this publication, provided study materials, reagents, materials, laboratory samples, instrumentation, computing resources, or other analysis tools. MEA, FV, managed and coordinated the research activity planning and execution. LC, MEA, JPS, YP, TB, conducted statistical, mathematical, computational, or other formal techniques to analyze or synthesize study data. JH, KF, RS, YP, PL, MEA, and BAR set up and conducted laboratory protocols and plant experiments. MEA, FV, JFS created genotype seed collections. The GrENE-net consortium contributed to experimental design and launched the GrENE-net.org experiments. LC, YP, JSP, PL, TB, MEA wrote the first manuscript draft, and all authors edited and reviewed the latest manuscript version.

## Supplemental Information Guide

Monitoring adaptation and demography of plant experimental populations with Pool-Sequencing

**<u>GrENE-net.org consortia authors</u>**

**<u>Supplemental Materials & Methods:</u> Extended DNA preparation, sequencing methods, and computational analyses.**

**<u>Supplemental Mathematical Appendix:</u> Population genetic equations adapted for Pool-Seq, including unbiased estimators of diversity and of F_ST_ for pool-sequencing data, and simulations.**

We provide this document online, see https://github.com/lczech/pool-seq-pop-gen-stats

## GrENE-net.org consortia authors

Delker, Carolin (1), Dimitrakopoulos, Panayiotis G. (2), Durka, Walter (3), Escribano-Avila, Gema (4), Franks, Steven J. (5), Fritschi, Felix B. (6), Galanidis, Alexandros (2), Garcia-Fernández, Alfredo (7), Hamann, Elena (8), Iriondo, J.M. (7), Juenger, Thomas E. (9), Lara-Romero, Carlos (7), Maag, Daniel (10), March-Salas, Martí (11), Morente-López, Javier (7), Pärtel, Meelis (12), Quint, Marcel (13), Seifan Merav (14), Snoek, Basten L. (15), Stam, Remco (16), Stinchcombe, John R. (17), Till-Bottraud, Irène (18), Traveset, Anna (19), Valay, Jean-Gabriel (20), Van Zanten, Martijn (21), Violle, Cyrille (22), Wódkiewicz Maciej (23)

(1) Crop Physiology, Institute of Agricultural and Nutritional Sciences, Martin Luther University Halle-Wittenberg, Germany

(2) Biodiversity Conservation Laboratory, Department of Environment, University of the Aegean, 81100 Mytilene, Lesbos, Greece

(3) Department of Community Ecology, Helmholtz Centre for Environmental Research-UFZ, 06120 Halle, Germany

(4) Arboriculture Department,Tecnigral SL, Madrid, Spain.

(5) Department of Biological Sciences and the Louis Calder Center, Fordham University, Bronx, New York, USA

(6) Division of Plant Science & Technology, University of Missouri, Columbia, MO, USA

(7) ECOEVO group, Biodiversity and Conservation Area, ESCET, Rey Juan Carlos University, Móstoles, Spain

(8) Biological Sciences, Fordham University, Bronx, NY, USA

(9) University of Texas at Austin, Department of Integrative Biology, Austin, USA

(10) Department of Pharmaceutical Biology, Julius-von-Sachs-Institute, Julius-Maximilians-Universität Würzburg, Würzburg, Germany

(11) Plant Evolutionary Ecology, Faculty of Biological Sciences, Goethe University Frankfurt, Max-von-Laue-Str. 13, 60438 Frankfurt am Main, Germany

(12) Institute of Ecology and Earth Sciences, University of Tartu,Tartu, Estonia

(13) Institute of Agricultural and Nutritional Sciences, Martin Luther University Halle-Wittenberg, Halle (Saale), Germany, German Centre for Integrative Biodiversity Research (iDiv), Halle-Jena-Leipzig, Germany

(14) Mitrani Department of Desert Ecology, Swiss Institute for Dryland Environmental & Energy Research, Blaustein Institutes for Desert Research, Ben-Gurion University of the Negev, Israel

(15) Theoretical Biology & Bioinformatics, Utrecht University, Padualaan 8, 3584 CH Utrecht the Netherlands

(16) Chair of Phytopathology,Technical University of Munich, Germany

(17) Université Clermont Auvergne, CNRS, GEOLAB, 63000 Clermont Ferrand, France

(18) Global change research group, Mediterranean Institute of Advanced Studies, Esporles, Mallorca, Balearic Islands, Spain

(19) Lautaret garden, Université Grenoble Alpes, CNRS, Grenoble, France

(20) Molecular Plant Physiology, Department of Environmental Biology, Utrecht University, Utrecht,The Netherlands

(21) CEFE, Univ Montpellier, CNRS, EPHE, IRD, France

(22) Faculty of Biology, Biological and Chemical Research Centre,University of Warsaw,Warsaw, Poland

(23) Ecology & Evolutionary Biology, University of Toronto,Toronto, Canada

## Supplemental Materials & Methods for

### Plant growth protocol

#### Experiment 1

The 231 ecotypes were bulked in growth chambers at 20°C under the long-day condition (16 hours light / 8 hours dark) in three locations. In the Max Planck Institute for Biology, Germany, the growth chambers at the University of Tübingen’s Institute of Evolutionary Ecology, Germany, and the CNRS Centre for Functional and Evolutionary Ecology in Montpellier, France.

#### Experiment 2

The Col-0 and RUM-20 plants were grown in growth chambers at 22°C under the long-day condition.

#### Experiment 3

We took one tube containing 0.1 g of the founder seed mix (∼5,000 seeds), bleach-sterilized it, washed it (20 min; 500ml solution, 10% bleach, 20% SDS) and submerged the seeds in 1% agar solution for 5 days at 4 °C in the dark. Seeds were planted in trays with soil (CL-P, Einheitserde Werkverband e.V., Sinntal-Altengronau Germany) in 25 trays, 1,000 pots, and germinants were thinned to one plant per pot. We watered abundantly, growing the plants at 16 °C for 13 days (long day conditions) before a 60 day vernalization at 4 °C (short day conditions). The vernalization approach aimed to avoid flowering time differences among our diverse genotypes. Subsequently, the trays were transferred to 20 °C under the long-day condition for flowering. Two weeks later, 50 pots were randomly chosen to sample leaves and flowers whenever the plants bloomed

#### Experiment 4

The genotypes planted are 451 natural accessions (**Dataset S1**), all mixed together, across three plots (about 2 seeds/cm^2^ in 1m^2^ plots). 20 siliques from two different parental individuals of all genotypes were pooled and sowed in three batches: one in November 2014, one in February 2015, one in March 2015. On March 2, 2016, before flowering, bulk soil was taken to germinate seeds in the growth chamber and collect flowers for sequencing to avoid any selection of genotypes. 50-101 flowers from the plants growing in the growth chamber were sampled to generate allele frequency data for time point 0 (**Table S5**). In the field, 50-100 flowers, 80-200 flowers, and 60-300 flowers were sampled on April 1 (time point 1), April 22 (time point 2), and May 6 (time point 3) respectively and used for sequencing.The numbers of flowers sampled per plot are outlined in **Table S5**.

### DNA extraction

#### Experiments 1, 3, and 4

The GrENE-net founder seed mix, containing 231 natural accessions (**Dataset S1**, Exp 1), was aliquoted into eight replicates according to the tissue input amount recommended by the Qiagen DNeasy Plant Mini kit (Hilden, Germany) (**Table S1**). Seed aliquots were suspended in 0.1% agar and kept at 4℃ in the dark for 9 to 11 days to initiate germination. Then, seed aliquots were centrifuged and the supernatant was removed. 0.5 mL of rock and 800 µL of lysis buffer AP1 from the DNeasy kit were added to the seed tubes. Tissue homogenization was carried out using the Quickprep adapter in a FastPrep-24 (MP Biomedicals, Irvine, CA, USA) with the following setting: 6.0 m/sec for 40 seconds. Each tube was homogenized for a total of 2 rounds. 8 µL of RNase (100 mg/mL) from the DNeasy kit was added to the seed homogenate. After a short vortex and a quick spin, the seed homogenate was incubated at 65℃ for 10 minutes. After the incubation, 185 µL of buffer P3 from the DNeasy kit was added to the seed lysate. The tube was inverted and incubated on ice for 5 minutes. The rest of the extraction followed the standard Qiagen DNeasy Plant Mini protocol. DNA was eluted in 100 µL of AE buffer.

The leaf subsamples and flower subsamples (Exp. 3) were extracted similarly with the DNeasy Plant Mini kit (Qiagen, Hilden, Germany) with modifications to the grinding step. Tissue samples and 5 ceramic beads were placed in a screw-cap tube and froze with liquid nitrogen. The homogenization was again carried out using FastPrep-24 (MP Biomedicals, Irvine, CA, USA) with a different setting: 4.0 m/sec for 15 seconds. Each tube was homogenized for a total of 2 rounds. The rest of the extraction followed the standard DNeasy protocol.

The field experiment samples (Exp. 4) of pools of flowers (Table S6) were processed as previously (Exp. 3).

#### Experiment 2

Due to the high cost of commercial kits such as the Qiagen DNeasy Plant Mini kit ($4.46 per isolation at listed price), we set up a cheaper plate-based DNA extraction protocol based on the widely used 2x CTAB protocol (Doyle and Doyle, 1987). All GrENE-net DNA extracts were isolated using this custom protocol from pooled flower samples collected from the 45 field sites. We partially replicated the leaf and flower comparison (see Experiment 3 in the main text) using our CTAB/chloroform protocol. In the case of flowers, two flowers of similar size were collected in the same tube prior to DNA extraction (n = 3). In the case of leaves, a leaf was independently extracted, but leaf extracts from two distinct ecotypes, Col-0 and RUM-20, were combined at similar DNA mass prior to library preparation (n = 3) (see Experiment 2 in the main text).

2- mercaptoethanol was added to the 2x CTAB buffer (1.4 M NaCl, 100 mM Tris pH 8.0, 20 mM EDTA pH 8.0, 2% w/v CTAB, 1% w/v PVP, ddH_2_O) to a final concentration (v/v) of 0.3%. The buffer was warmed at 65°C for at least 30 minutes. Using a TissueLyser II (Qiagen, Hilden, Germany), frozen plant tissues were pulverized with 3.2 mm steel beads in 2.0 mL tubes on chilled adapter sets 2 x 24. Homogenization was carried out at 22/s for 35 sec and repeated until the frozen tissues attained the appearance of greenish white powders. 500 µL of pre-warmed 2x CTAB buffer was added to each tube to thoroughly resuspend the pulverized tissue. Samples were incubated at 65°C for 50 minutes and inverted every 10 to 15 minutes to resuspend the precipitates. After incubation, the lysate was transferred to a new 2.0 mL tube. When the lysate was cooled to room temperature, 500 µL of chloroform:isoamyl alcohol (24:1) was added to the lysate. The tube was vigorously shaken until the lysate and chloroform appeared well-mixed. The sample was centrifuged at 20,000 rcf for 14 minutes or until the upper aqueous layer appeared clear. 300 µL of the aqueous layer was transferred to a new tube or a 96-well deep well plate if doing high-throughput processing. 225 µL (0.75 vol) of isopropanol was added to the supernatant and mixed well by pipetting. The sample was incubated at 4°C for at least 30 minutes or at -20°C overnight. After incubation, the sample was centrifuged at max speed for 15 minutes in a tube. Alternatively, the 96-well plate was centrifuged at 6,100 rcf for 45 minutes. After discarding the supernatant, freshly prepared 70% ethanol was added to wash the DNA pellet. The sample was centrifuged at max speed for 5 minutes in a tube. Alternatively, the 96-well plate was centrifuged at 6,100 rcf for 30 minutes. The ethanol was removed and the pellet was left to air dry for 10 to 15 minutes. The DNA pellet was eluted in Tris buffer containing RNase A (10 mM Tris-HCl pH9.0, ddH_2_O, 20 µg/mL RNase A). The eluate was incubated at 37°C for 30 minutes. After a pulse spin, the DNA extract was stored at -20°C.

### Library preparation

#### Experiment 1

Because the total number of samples was small and we had large amounts of DNA (**Table S1**), we conducted library preparations with Illumina’s TruSeq PCR free library kit (Ilumina, San Diego, California).

#### Experiment 2

Because there were only 11 samples, tube-based quantification was performed using the Qubit dsDNA HS assay. The readings for the input DNA concentration are documented in Table S2. The library preparation protocol was based on (Baym et al., 2015) with some modifications. 2 µL of DNA sample was mixed with 2.75 µL TD buffer (Tagment DNA Buffer) and 0.25 µL TD enzyme (Mira Loma, California, USA). The tagmentation reaction mixture was mixed well by gentle pipetting. After a flash spin, the sample was incubated at 55°C for 10 minutes and held at 10°C.

Once equilibrated to room temperature or lower, the samples were flash spinned. Then, the tagmented DNA was mixed with 8 µL 2x KAPA HiFi HotStart ReadyMix (KAPA Biosystems, Boston, MA, USA), 1.5 µL 10 µM P5 indexing primers (final concentration 0.75 µM), 1.5 µL 10 µM P7 indexing primers (equimolar to P5), and 4 µL Tris-Cl buffer (pH 8.0). The PCR reaction mixture was mixed well by gentle pipetting and the liquid was spinned down. The DNA was amplified using the following thermal cycling program:

1. 72°C for 3 minutes
2. 95°C for 3 minutes
3. 98°C for 20 seconds
4. 63°C for 30 seconds
5. 72°C for 30 seconds
6. Repeat from step 3 for 11 additional cycles (i.e. a total of 12 cycles)
7. 72°C for 5 minutes
8. 10°C hold

For post-amplification cleanup and size selection, the 11 libraries were multiplexed in a 1.5 mL tube by mixing 10 µL of each library. The library volume was estimated by aspirating with a P200 pipette (V_lib_). 0.45 volume (i.e. 0.45 x V_lib_) of homemade SPRI beads was added to the 11-plex library. The tube was incubated for 5 minutes on a regular rack and then incubated for 5 minutes on a magnet stand until a bead pellet forms. This bead pellet represents the first elution fraction. The supernatant excluding the first pellet was transferred to a new 1.5 mL tube on a regular rack. 0.6 volume (i.e. [0.6 - 0.45] x V_lib_) of homemade SPRI beads was added to the supernatant. The tube was incubated for 5 minutes on a regular rack and then incubated for 5 minutes on a magnet stand until a bead pellet forms. This bead pellet represents the second elution fraction. The supernatant excluding the second pellet was removed. Each magnetic bead pellet was washed by gently adding 700 µL 70% ethanol and was incubated for 30 seconds before removing the ethanol. The ethanol wash was repeated once. The bead pellet was air dried until they lost the shine and began showing tiny cracks.

The tube containing the bead pellet was taken off the magnet and resuspended in 36 µL of AE buffer (10 mM Tris-Cl, 0.5 mM EDTA; pH 9.0). After incubating on a regular rack for 3 minutes, the tube was put on magnet stands for 5 minutes until the bead pellet formed. 34 µL of the eluate fraction was transferred to a new 1.5 mL tube. Both eluted fractions were quantified with Qubit and analyzed on a TapeStation 4150 (Memphis, Tennessee, USA) using a D1000 ScreenTape (Cedar Creek, Texas, USA).The second fraction was sequenced on a HiSeq 2 x 150 lane ( **Fig. S6** ).

#### Experiment 3

To compare seeds with flowers and leaf extracts without library preparation differences, we conducted library preparations with Illumina’s TruSeq PCR library kit (Ilumina, San Diego, California) as in Experiment 1.

#### Experiment 4

The library preparation procedure was similar to what was described in Rowan et al. 2019 *Genetics* with minor volume adjustments. Specifically, the twelve amplified libraries were multiplexed together by mixing 5 µL of each library. The total volume was brought up to 100 µL with 10 mM Tris-Cl (pH 8.5). The rest of the size selection was performed as written in Rowan et al. (2019). The fragment length distribution of bead fraction 3 was verified with a Bioanalyzer before being sent for sequencing on an Illumina HiSeq 3000.

## Supplemental Datasets and Tables

**Dataset S1 | Ecotype IDs used for outdoor experiment and GrENE-net seed mixture**

Metadata of the 451 and 231 ecotypes lists.

[<google drive link>](https://docs.google.com/spreadsheets/d/1fdmzxI-bCipHfAUOEo55ipFVeQc54oVmwv8ANTS3dak/edit?usp=sharing)

**Dataset S2 | Genes within 10Kb regions with F_ST_ > 0.2 in 3+ E&R replicates**

TAIR summary of gene annotation.

[<google drive link>](https://docs.google.com/spreadsheets/d/1jqs4aS1Ds2qTVIXKnQEIwY5oPC1Z1gL7NdAQkhI0rRY/edit?usp=sharing)

**Table S1.** GrENE-net founder seed mix DNA extraction replicates.

| Sample identifier | Tissue amount (mg) | Approximate number of individuals | DNA concentration (ng/μL) |
| --- | --- | --- | --- |
| GrENE-net 231 founder seed mix #1 | 100 | 5000 | 7.98 |
| #2 | 100 | 5000 | 6.48 |
| #3 | 100 | 5000 | 8.56 |
| #4 | 17.4 | 870 | 21.8 |
| #5 | 18 | 900 | 24.8 |
| #6 | 18.3 | 915 | 23.6 |
| #7 | 20 | 1000 | 24.4 |
| #8 | 21.6 | 1080 | 25 |

**Table S2.** Input DNA concentration library preparation for validation experiments.

| Sample identifier | Input concentration (ng/μL) |
| --- | --- |
| Col-0 leaf extract | 2.17 |
| RUM-20 leaf extract | 2.04 |
| Pooled flower extract #1 | 1.84 |
| Pooled flower extract #3 | 1.97 |
| Pooled flower extract #5 | 1.98 |
| Col-RUM leaf extract pool #1 | 2 (predicted, 2 μL of 8.33 ng/μL diluted in 6.33 μL Tris buffer) |
| Col-RUM leaf extract pool #2 | 2 (predicted, 2 μL of 8.53 ng/μL diluted in 6.53 μL Tris buffer) |
| Col-RUM leaf extract pool #3 | 2 (predicted, 2 μL of 8.71 ng/μL diluted in 6.71 μL Tris buffer) |

\*due to the flexibility (i.e. $\leq 3$ ng/ μL) in the acceptable input range for this protocol, the three leaf extract pools were not quantified after dilution

**Table S3.**
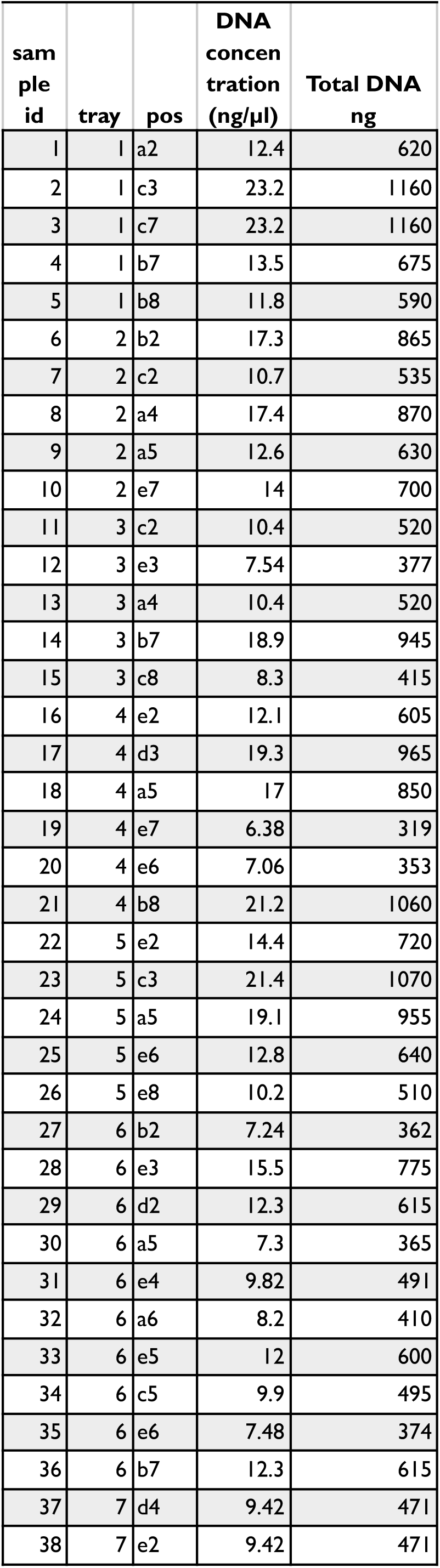
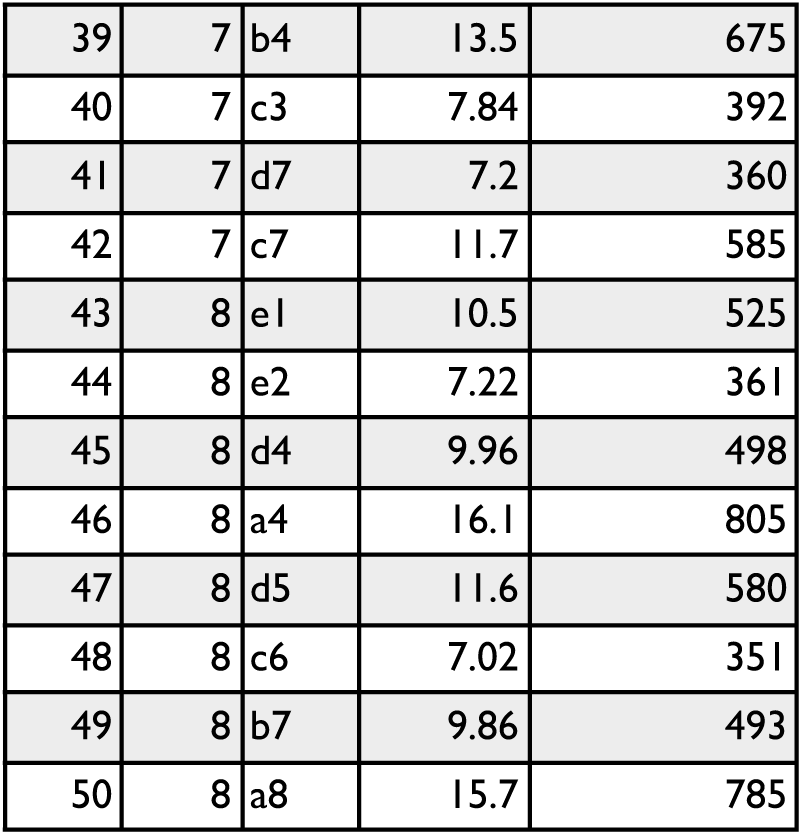
Sampling of 50 leaves.

| sample id | tray | pos | DNA concentration (ng/μl) | Total DNA ng |
| --- | --- | --- | --- | --- |
| 1 | 1 | a2 | 12.4 | 620 |
| 2 | 1 | c3 | 23.2 | 1160 |
| 3 | 1 | c7 | 23.2 | 1160 |
| 4 | 1 | b7 | 13.5 | 675 |
| 5 | 1 | b8 | 11.8 | 590 |
| 6 | 2 | b2 | 17.3 | 865 |
| 7 | 2 | c2 | 10.7 | 535 |
| 8 | 2 | a4 | 17.4 | 870 |
| 9 | 2 | a5 | 12.6 | 630 |
| 10 | 2 | e7 | 14 | 700 |
| 11 | 3 | c2 | 10.4 | 520 |
| 12 | 3 | e3 | 7.54 | 377 |
| 13 | 3 | a4 | 10.4 | 520 |
| 14 | 3 | b7 | 18.9 | 945 |
| 15 | 3 | c8 | 8.3 | 415 |
| 16 | 4 | e2 | 12.1 | 605 |
| 17 | 4 | d3 | 19.3 | 965 |
| 18 | 4 | a5 | 17 | 850 |
| 19 | 4 | e7 | 6.38 | 319 |
| 20 | 4 | e6 | 7.06 | 353 |
| 21 | 4 | b8 | 21.2 | 1060 |
| 22 | 5 | e2 | 14.4 | 720 |
| 23 | 5 | c3 | 21.4 | 1070 |
| 24 | 5 | a5 | 19.1 | 955 |
| 25 | 5 | e6 | 12.8 | 640 |
| 26 | 5 | e8 | 10.2 | 510 |
| 27 | 6 | b2 | 7.24 | 362 |
| 28 | 6 | e3 | 15.5 | 775 |
| 29 | 6 | d2 | 12.3 | 615 |
| 30 | 6 | a5 | 7.3 | 365 |
| 31 | 6 | e4 | 9.82 | 491 |
| 32 | 6 | a6 | 8.2 | 410 |
| 33 | 6 | e5 | 12 | 600 |
| 34 | 6 | c5 | 9.9 | 495 |
| 35 | 6 | e6 | 7.48 | 374 |
| 36 | 6 | b7 | 12.3 | 615 |
| 37 | 7 | d4 | 9.42 | 471 |
| 38 | 7 | e2 | 9.42 | 471 |
| 39 | 7 | b4 | 13.5 | 675 |
| 40 | 7 | c3 | 7.84 | 392 |
| 41 | 7 | d7 | 7.2 | 360 |
| 42 | 7 | c7 | 11.7 | 585 |
| 43 | 8 | e1 | 10.5 | 525 |
| 44 | 8 | e2 | 7.22 | 361 |
| 45 | 8 | d4 | 9.96 | 498 |
| 46 | 8 | a4 | 16.1 | 805 |
| 47 | 8 | d5 | 11.6 | 580 |
| 48 | 8 | c6 | 7.02 | 351 |
| 49 | 8 | b7 | 9.86 | 493 |
| 50 | 8 | a8 | 15.7 | 785 |

**Table S4.** Combinatorics of flower and leaf pooling. The 50 randomly selected plants were sampled for the 50 and 100 samples as well as for nested samples of smaller sets of plants. A graphical scheme of this sampling is in **Fig. S3** .

| sample id | tray | pos | x100 | x50a | x50b | x25a | x25b | x10b1 | x10b2 | x5 |
| --- | --- | --- | --- | --- | --- | --- | --- | --- | --- | --- |
| 1 | 1 | a2 | yes | yes | yes |  | yes |  |  | yes |
| 2 | 1 | c3 | yes | yes | yes |  | yes | yes |  |  |
| 3 | 1 | c7 | yes | yes | yes | yes |  |  |  |  |
| 4 | 1 | b7 | yes | yes | yes |  | yes |  | yes |  |
| 5 | 1 | b8 | yes | yes | yes | yes |  |  |  |  |
| 6 | 2 | b2 | yes | yes | yes |  | yes | yes |  |  |
| 7 | 2 | c2 | yes | yes | yes |  | yes | yes |  |  |
| 8 | 2 | a4 | yes | yes | yes |  | yes | yes |  |  |
| 9 | 2 | a5 | yes | yes | yes | yes |  |  |  |  |
| 10 | 2 | e7 | yes | yes | yes |  | yes |  | yes |  |
| 11 | 3 | c2 | yes | yes | yes |  | yes | yes |  |  |
| 12 | 3 | e3 | yes | yes | yes |  | yes |  |  | yes |
| 13 | 3 | a4 | yes | yes | yes |  | yes | yes |  |  |
| 14 | 3 | b7 | yes | yes | yes | yes |  |  |  |  |
| 15 | 3 | c8 | yes | yes | yes |  | yes | yes |  |  |
| 16 | 4 | e2 | yes | yes | yes | yes |  |  |  |  |
| 17 | 4 | d3 | yes | yes | yes | yes |  |  |  |  |
| 18 | 4 | a5 | yes | yes | yes |  | yes |  | yes |  |
| 19 | 4 | e7 | yes | yes | yes | yes |  |  |  |  |
| 20 | 4 | e6 | yes | yes | yes |  | yes |  | yes |  |
| 21 | 4 | b8 | yes | yes | yes | yes |  |  |  |  |
| 22 | 5 | e2 | yes | yes | yes |  | yes | yes |  |  |
| 23 | 5 | c3 | yes | yes | yes | yes |  |  |  |  |
| 24 | 5 | a5 | yes | yes | yes |  | yes |  | yes |  |
| 25 | 5 | e6 | yes | yes | yes | yes |  |  |  |  |
| 26 | 5 | e8 | yes | yes | yes | yes |  |  |  |  |
| 27 | 6 | b2 | yes | yes | yes | yes |  |  |  |  |
| 28 | 6 | e3 | yes | yes | yes | yes |  |  |  |  |
| 29 | 6 | d2 | yes | yes | yes |  | yes |  |  | yes |
| 30 | 6 | a5 | yes | yes | yes | yes |  |  |  |  |
| 31 | 6 | e4 | yes | yes | yes | yes |  |  |  |  |
| 32 | 6 | a6 | yes | yes | yes | yes |  |  |  |  |
| 33 | 6 | e5 | yes | yes | yes |  | yes |  |  | yes |
| 34 | 6 | c5 | yes | yes | yes | yes |  |  |  |  |
| 35 | 6 | e6 | yes | yes | yes | yes |  |  |  |  |
| 36 | 6 | b7 | yes | yes | yes | yes |  |  |  |  |
| 37 | 7 | d4 | yes | yes | yes | yes |  |  |  |  |
| 38 | 7 | e2 | yes | yes | yes |  | yes |  | yes |  |
| 39 | 7 | b4 | yes | yes | yes |  | yes | yes |  |  |
| 40 | 7 | c3 | yes | yes | yes |  | yes |  | yes |  |
| 41 | 7 | d7 | yes | yes | yes |  | yes |  | yes |  |
| 42 | 7 | c7 | yes | yes | yes | yes |  |  |  |  |
| 43 | 8 | e1 | yes | yes | yes |  | yes |  | yes |  |
| 44 | 8 | e2 | yes | yes | yes | yes |  |  |  |  |
| 45 | 8 | d4 | yes | yes | yes |  | yes |  | yes |  |
| 46 | 8 | a4 | yes | yes | yes | yes |  |  |  |  |
| 47 | 8 | d5 | yes | yes | yes | yes |  |  |  |  |
| 48 | 8 | c6 | yes | yes | yes | yes |  |  |  |  |
| 49 | 8 | b7 | yes | yes | yes |  | yes | yes |  |  |
| 50 | 8 | a8 | yes | yes | yes |  | yes |  |  | yes |

**Table S5.** Pool extraction of flower combinatorics.

| Tube ID | DNA<br>concentrati<br>on (ng/ul) |
| --- | --- |
| 100 I | 20.8 |
| 100 2 | 20.8 |
| 50 A | 31.8 |
| 50 B | 33.2 |
| 25 A | 23.8 |
| 25 B | 19.5 |
| 10 B2 | 5.48 |
| 5 B | 4.68 |

**Table S6.** Sampling of pilot field experiment (Exp. 4) A total of 12 samples were sequenced either from seed banks or flowers of surviving plants in the outdoor field. Sequencing metrics are provided for each sample and the final working in silico pool.

| S | Time | rep | ID | # flowers | origin | Input DNA |  | bp | coverage | pool coverage |
| --- | --- | --- | --- | --- | --- | --- | --- | --- | --- | --- |
|  |  |  |  |  |  | (ng/μL) | read count |  | e |  |
| 1 | 0 | 1 | 1_0 | 56 | Seed bank | 0.376 | 40214556 | 6032183400 | 48.3 | 199.6 |
| 2 | 0 | 2 | 2_0 | 69 | Seed bank | 0.366 | 62882228 | 9432334200 | 75.5 |  |
| 3 | 0 | 3 | 3_0 | 101 | Seed bank | 0.424 | 63206340 | 9480951000 | 75.8 |  |
| 4 | 1 | 1 | 1_1 | 80 | Flowers from field | 0.404 | 51991084 | 7798662600 | 62.4 | 219.3 |
| 5 | 1 | 2 | 2_1 | 160 | Flowers from field | 0.474 | 65024068 | 9753610200 | 78 |  |
| 6 | 1 | 3 | 3_1 | 200 | Flowers from field | 0.454 | 65731096 | 9859664400 | 78.9 |  |
| 7 | 2 | 1 | 1_2 | 65 | Flowers from field | 0.374 | 71196816 | 1.068E+10 | 85.4 | 206.3 |
| 8 | 2 | 2 | 2_2 | 205 | Flowers from field | 0.452 | 47974450 | 7196167500 | 57.6 |  |
| 9 | 2 | 3 | 3_2 | 296 | Flowers from field | 0.33 | 52761222 | 7914183300 | 63.3 |  |
| 10 | 3 | 1 | 1_3 | 19 | Flowers from field | 0.25 | 60332690 | 9049903500 | 72.4 | 206.3 |
| 11 | 3 | 2 | 2_3 | 50 | Flowers from field | 0.434 | 102265250 | 15339787500 | 122.7 |  |
| 12 | 3 | 3 | 3_3 | 97 | Flowers from field | 0.452 | 64052368 | 9607855200 | 76.9 |  |

## Supplemental Figures

**Fig. S1.**
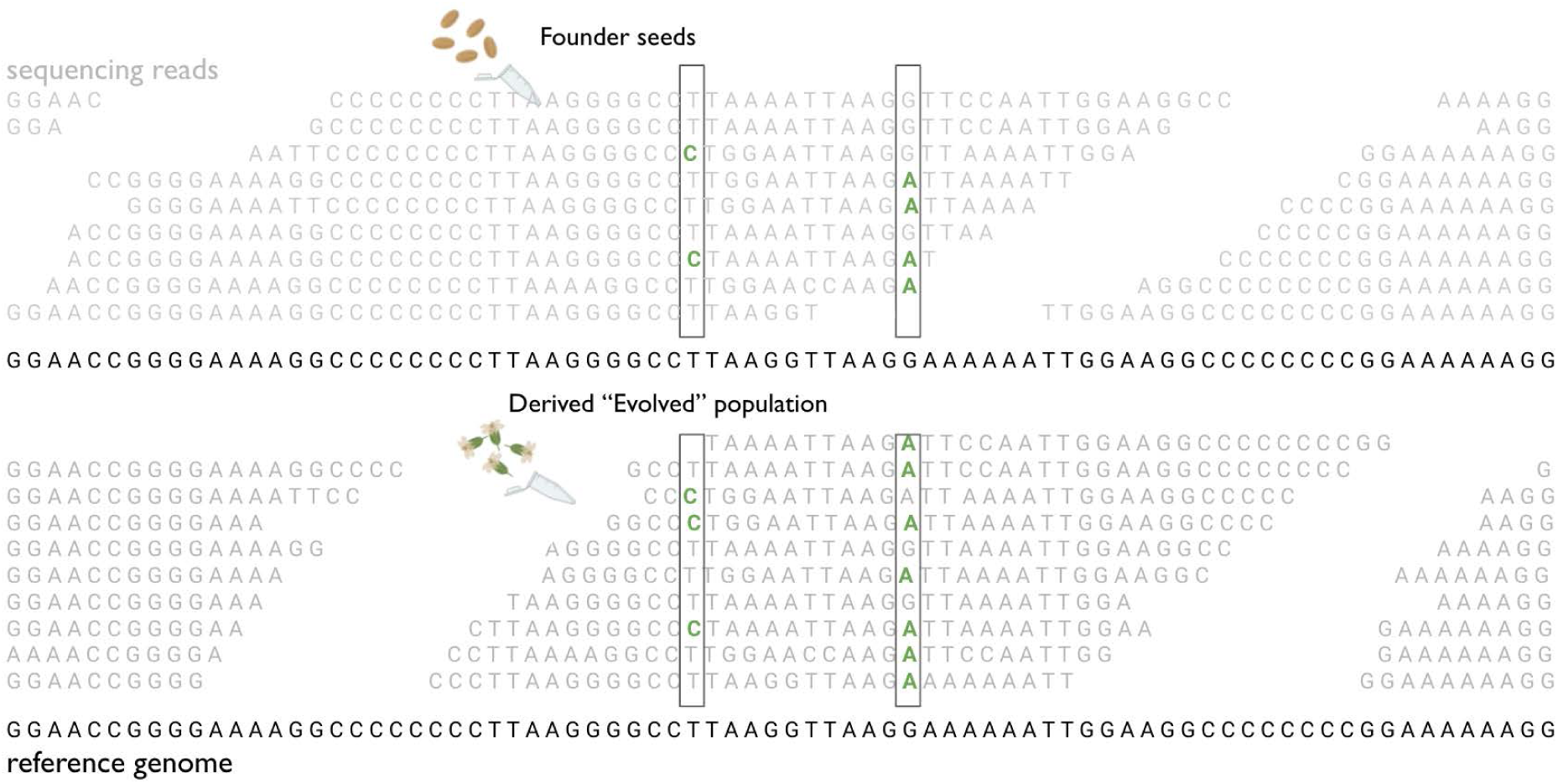
Cartoon of rationale of Pool-Seq. Reads from Illumina sequencing (typically ∼150 bp, not at scale) “piled” against the region of the genome where they map to. The rationale is that if founder allele frequencies, or a reference sample that did not experience natural selection, we can extract meaningful evolutionary insights from comparing those with “evolved” populations, i.e. those that have grown in outdoor environments. However, in the Pool Sequencing context, we have to deal with two sources of noise from a nested Binomial sampling: First, only a finite number of individuals is sampled from the population, and second, only a finite number of reads are sampled (i.e., processed) during sequencing.

**Fig. S2.**
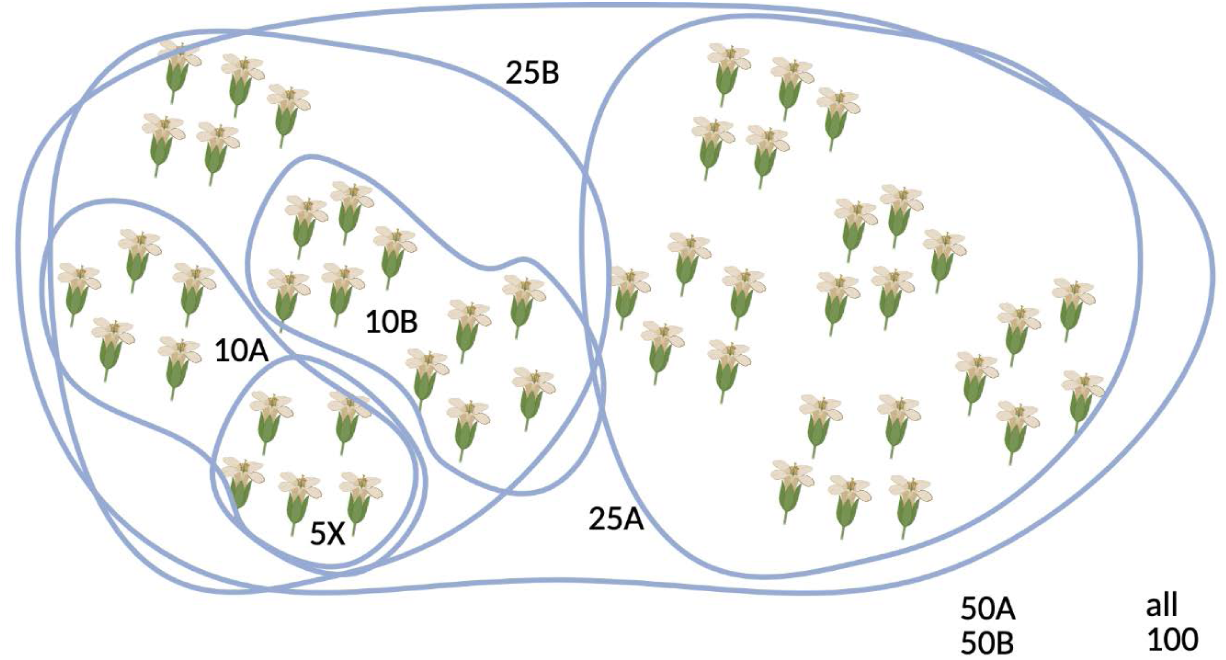
Hierarchical sampling of flowers for Pool-sequencing of different sizes. The selection of 50 individuals of Experiment 3 was conducted randomly from ∼2,500 plants, but smaller sets of individuals were conducted in a nested fashion (e.g. the 25B samples were the same individuals as the 10A, 10B, and 5X sample. This may enable downstream allele frequency comparisons).

**Fig. S3.**
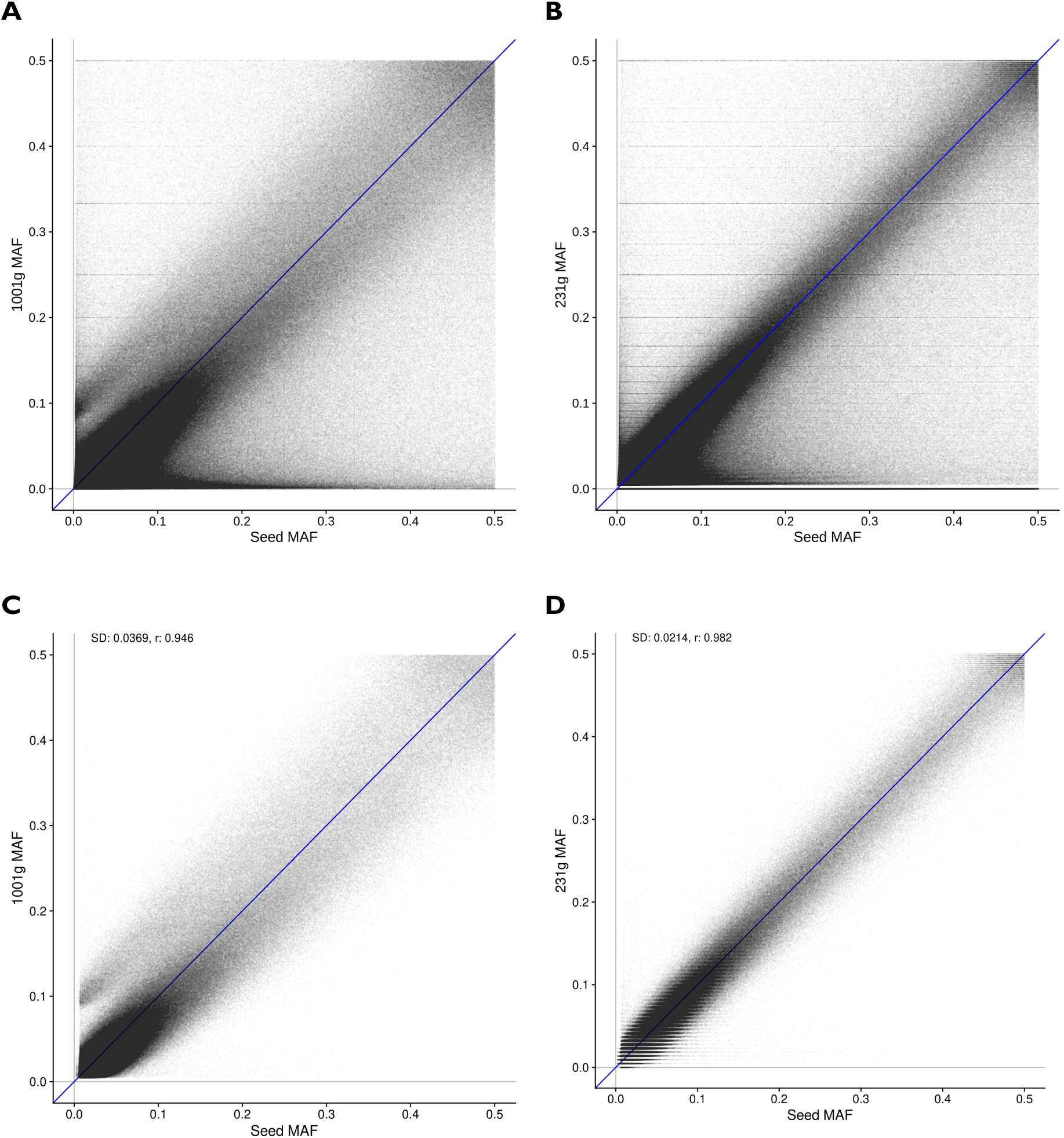
Allele frequencies from the 1001 Arabidopsis Genomes and from seed Pool-seq. The x-axis is the folded seed founder frequency used to source outdoor experiments of GrENE-net.org, based on counting nucleotides at each locus in a bam/pileup file of the mapped reads (we converted bam to pileup for easier file parsing; same in all bam-based plots below). The y-axis is the folded frequency characterized from (**A**) the 1001 Genomes VCF and (**B**) the GrENE-net founder VCF. (**C**) and (**D**) are the same comparisons as (A) and (B) but for only *bona fide* of 1,353,386 biallelic SNPs from the 515g subset of the 1001 Genomes.

**Fig. S4.**
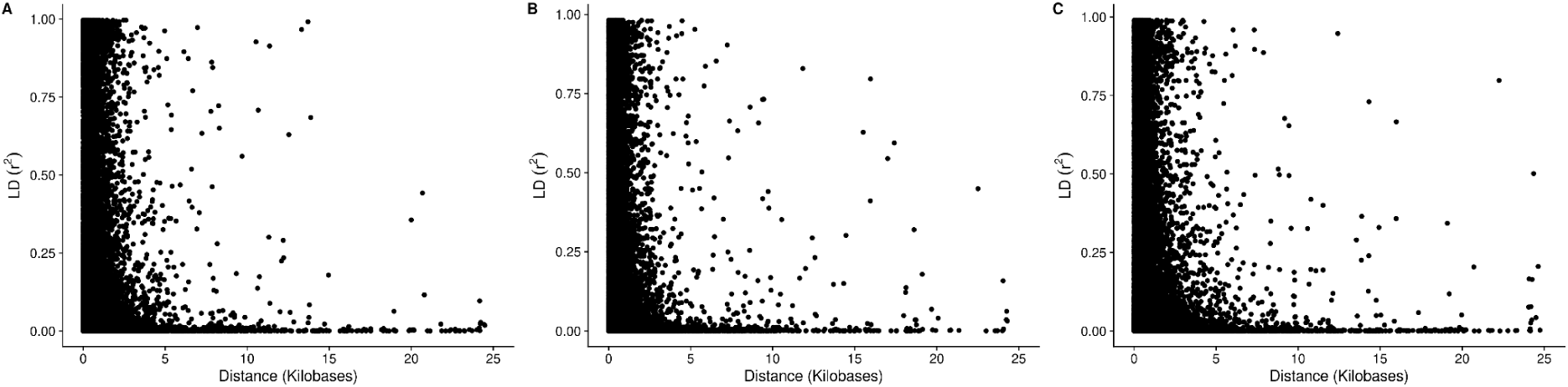
LD decay in the 1001G, the 470 and 231 and accessions sets. Linkage Disequilibrium LD decay using r^2^ for the genome collection of (**A**) the 1001 Genomes Project, (**B**) a subset of 231 used in Experiment 1 and 3, and (**C**) a subset of 451 of the 1001 Genomes used in Experiment 4.

**Fig. S5.**
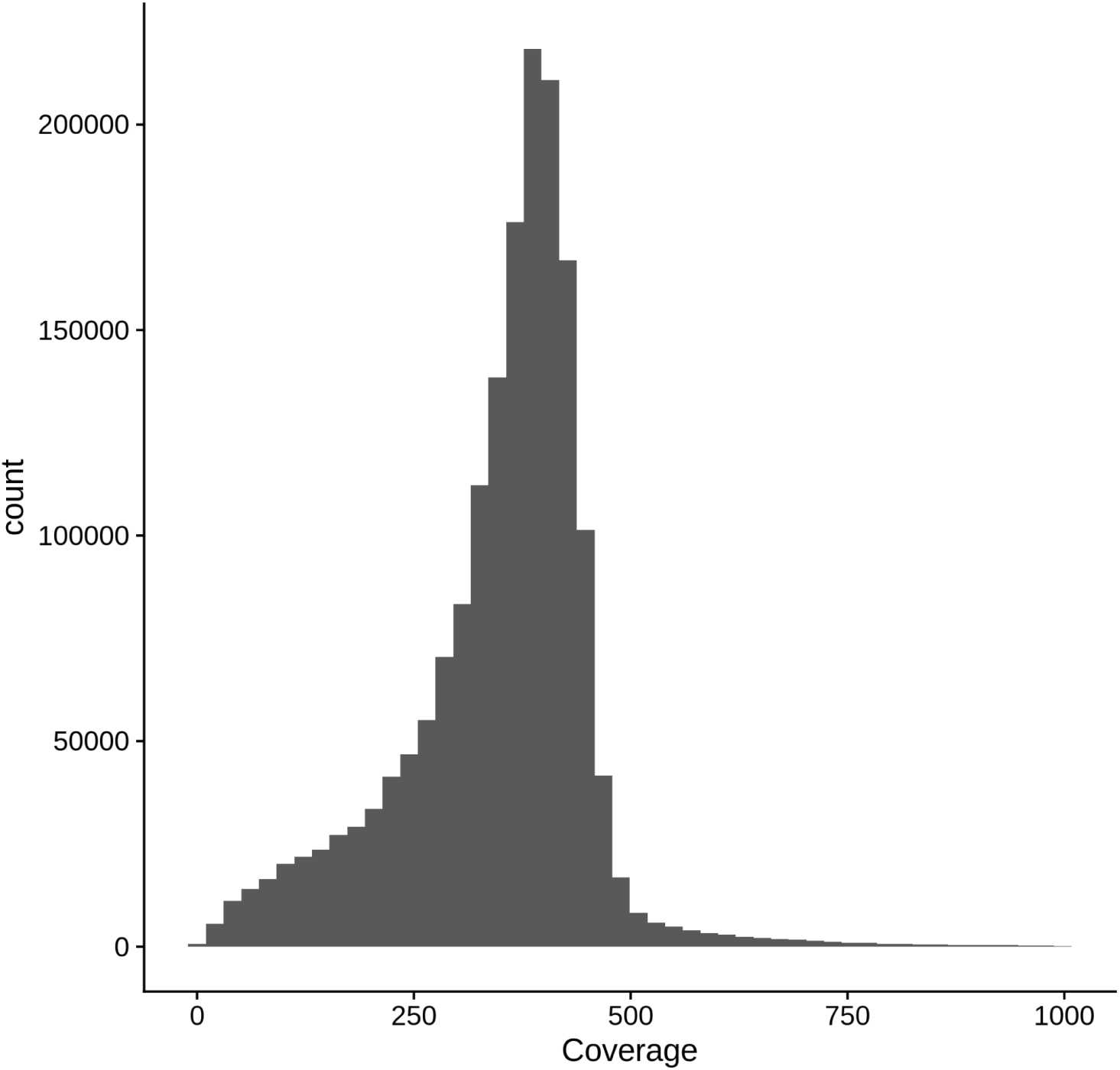
Coverage of the seed sequencing. Example of the coverage of the seeds as a *in silico* merge of 8 library samples (see **Table S1**) used in

**Fig. S6.**
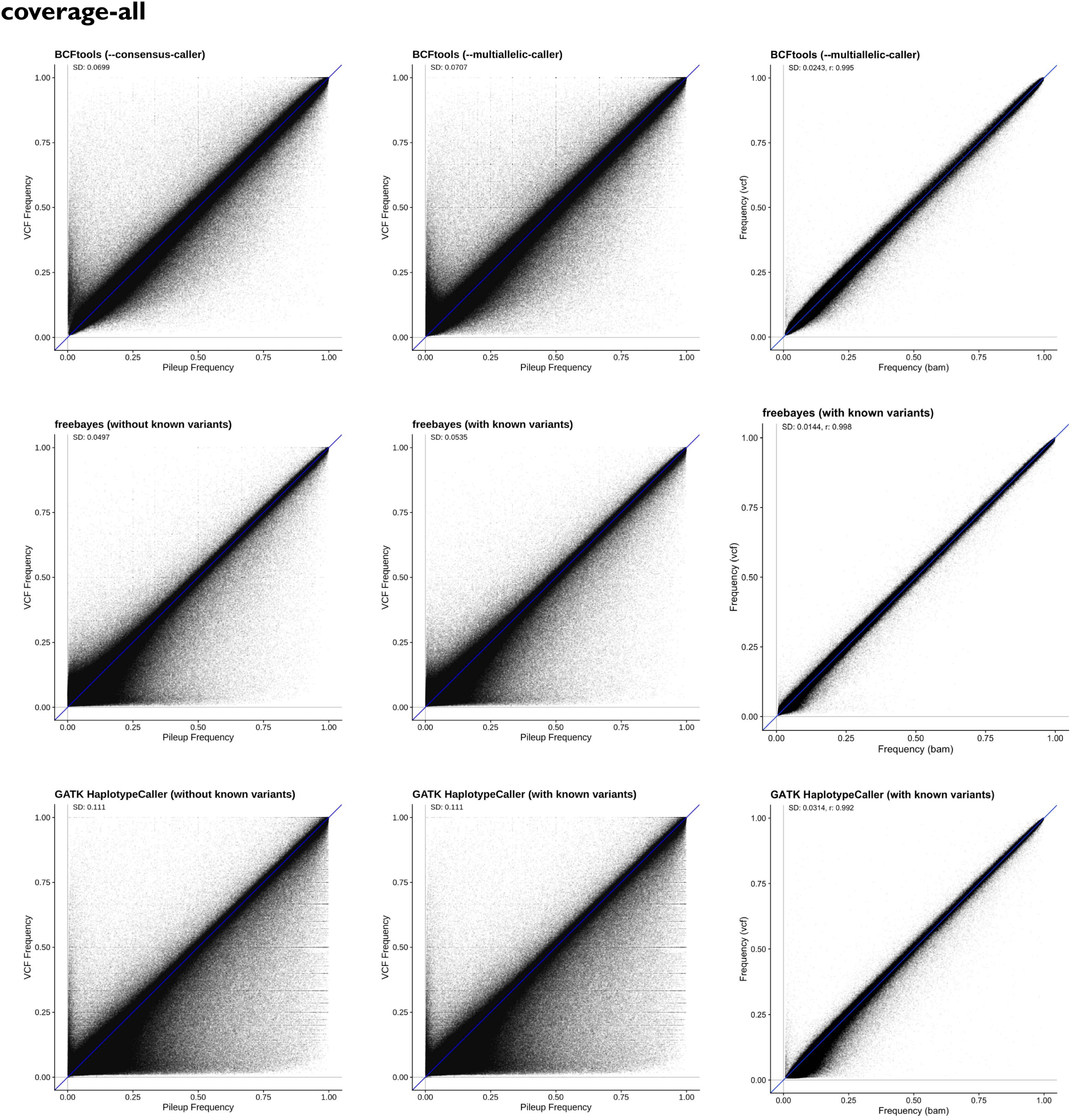
Correlation between raw frequencies and SNP caller fields (all coverages) The x-axis shows the allele frequency of a biallelic SNP based on bam/pileup format where raw ratios of alternative and reference bases are computed, and is shared across all comparisons. The y-axis shows the allele frequency of the same biallelic SNPs as inferred from the allelic depth (“AD”) VCF field from SNP calling outputs. Deviations from the y=x axis likely reveal artifacts created by SNP calling softwares. Each row presents three typical callers: BCFtools, freebayes, and GATK. Each column represents different calling mode or filterings: ‘free’ or discovery SNP calling, guided SNP calling at known variable positions from the 1001 Genomes, and subset of SNPs to *bona fide* of 1,353,386 biallelic SNPs from the 515g subset of the 1001 Genomes.

**Fig. S7.**
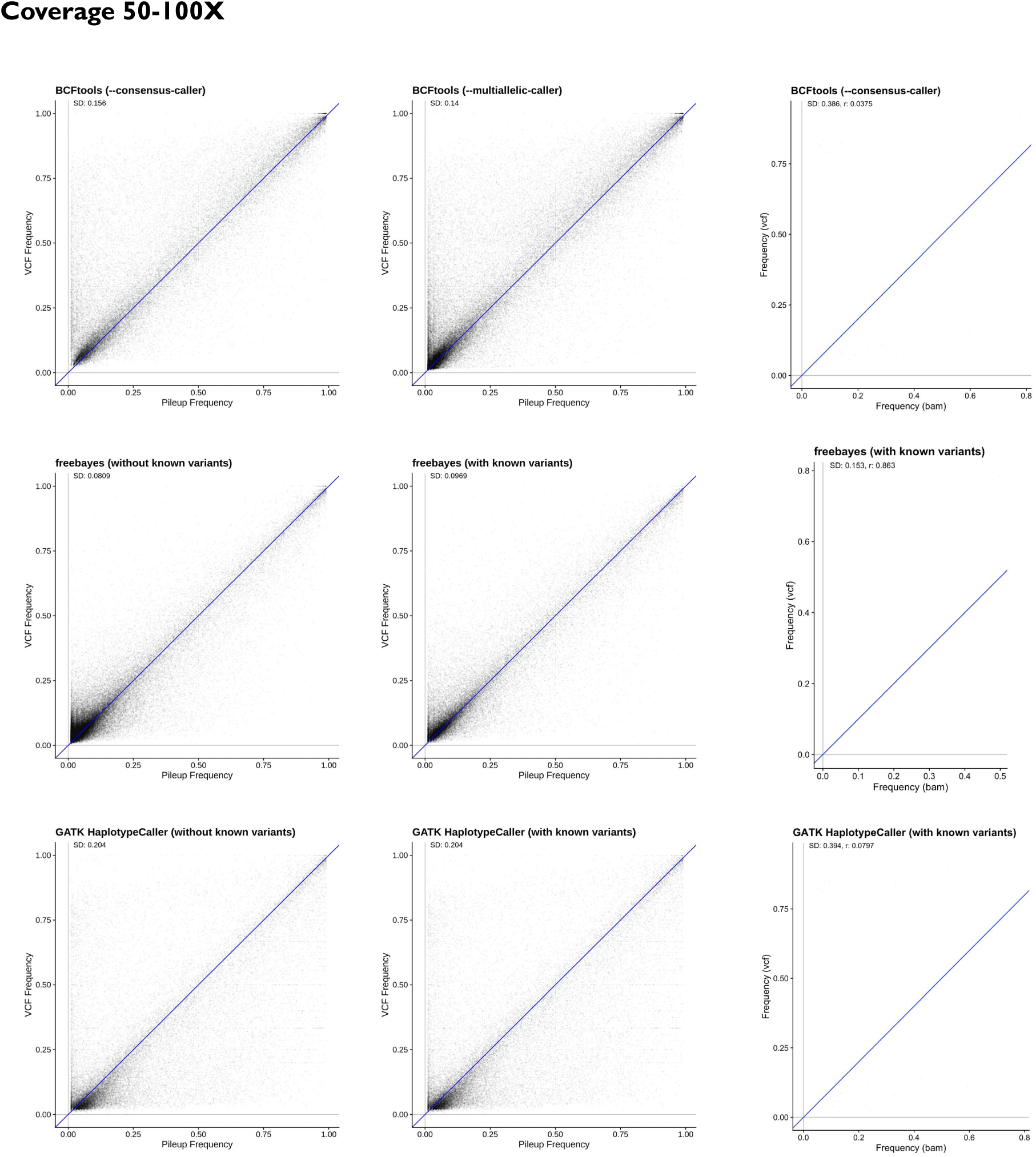
Correlation between raw frequencies and SNP caller fields (50-100X) Same as **Fig. S6** , but only for coverage 50-100X.

**Fig. S8.**
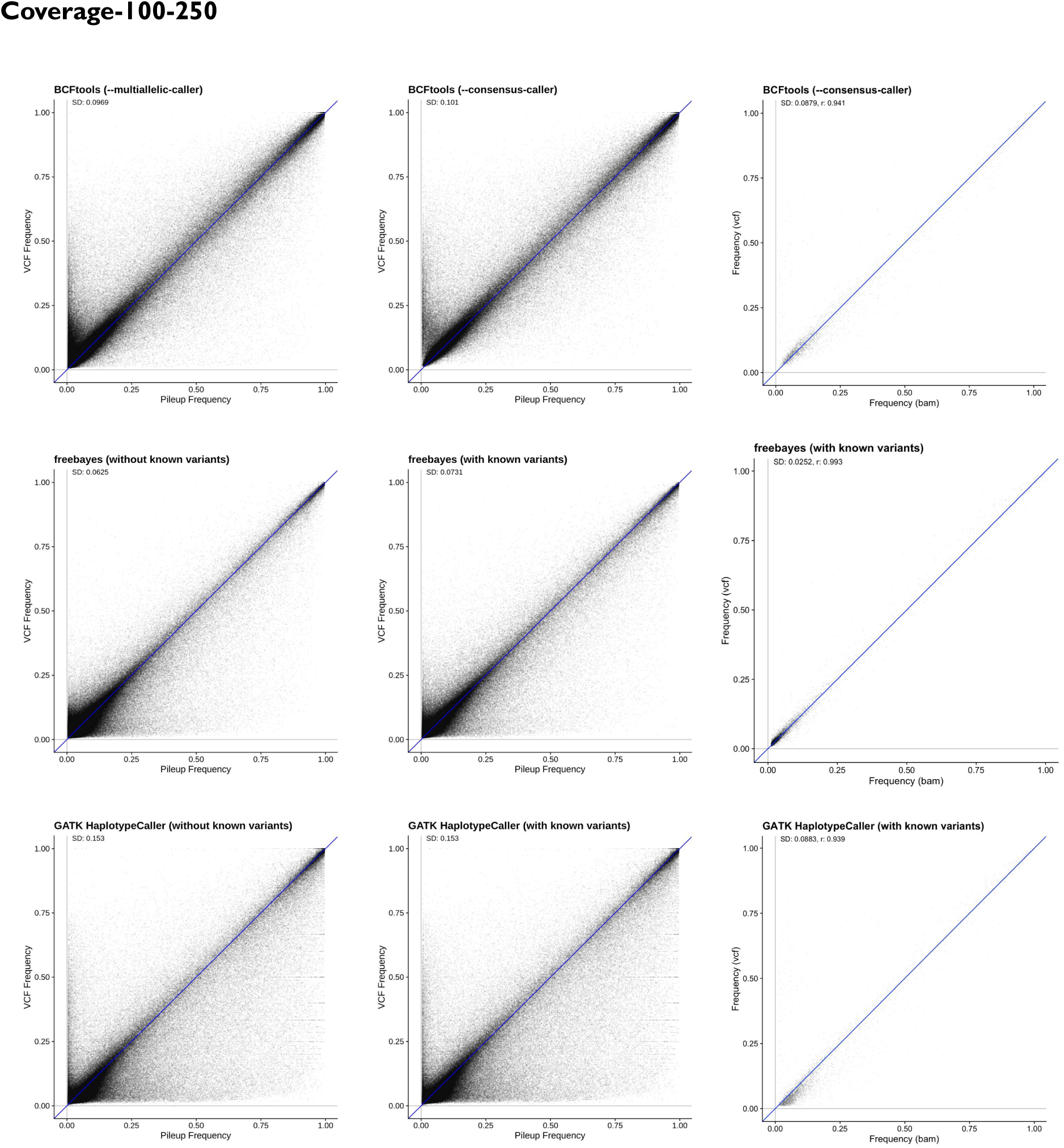
Correlation between raw frequencies and SNP caller fields (100-250X) Same as **Fig. S6** , but only for coverage 100-250X.

**Fig. S9.**
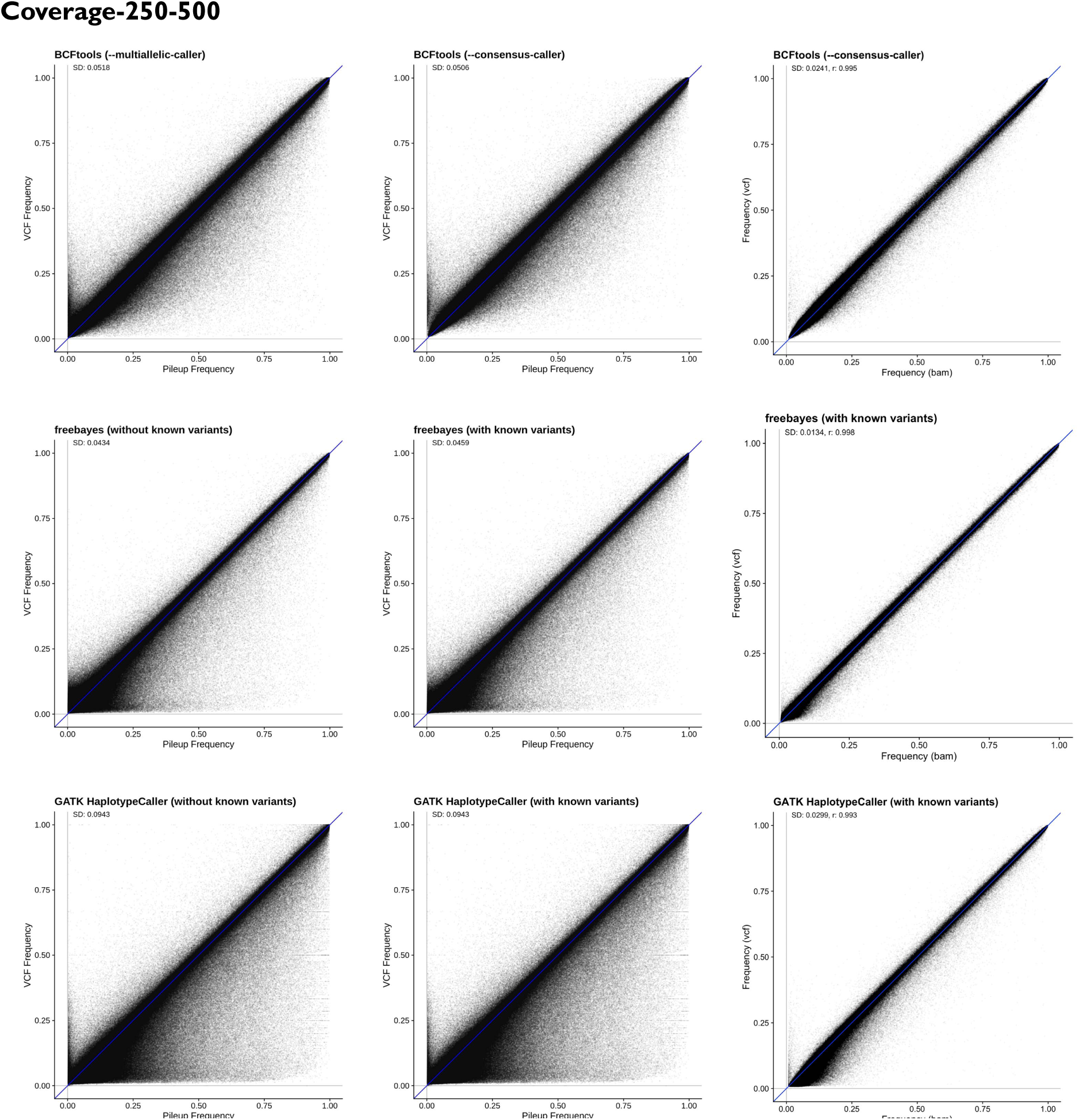
Correlation between raw frequencies and SNP caller fields (250-500X) Same as **Fig. S6** , but only for coverage 250-500X.

**Fig. S10.**
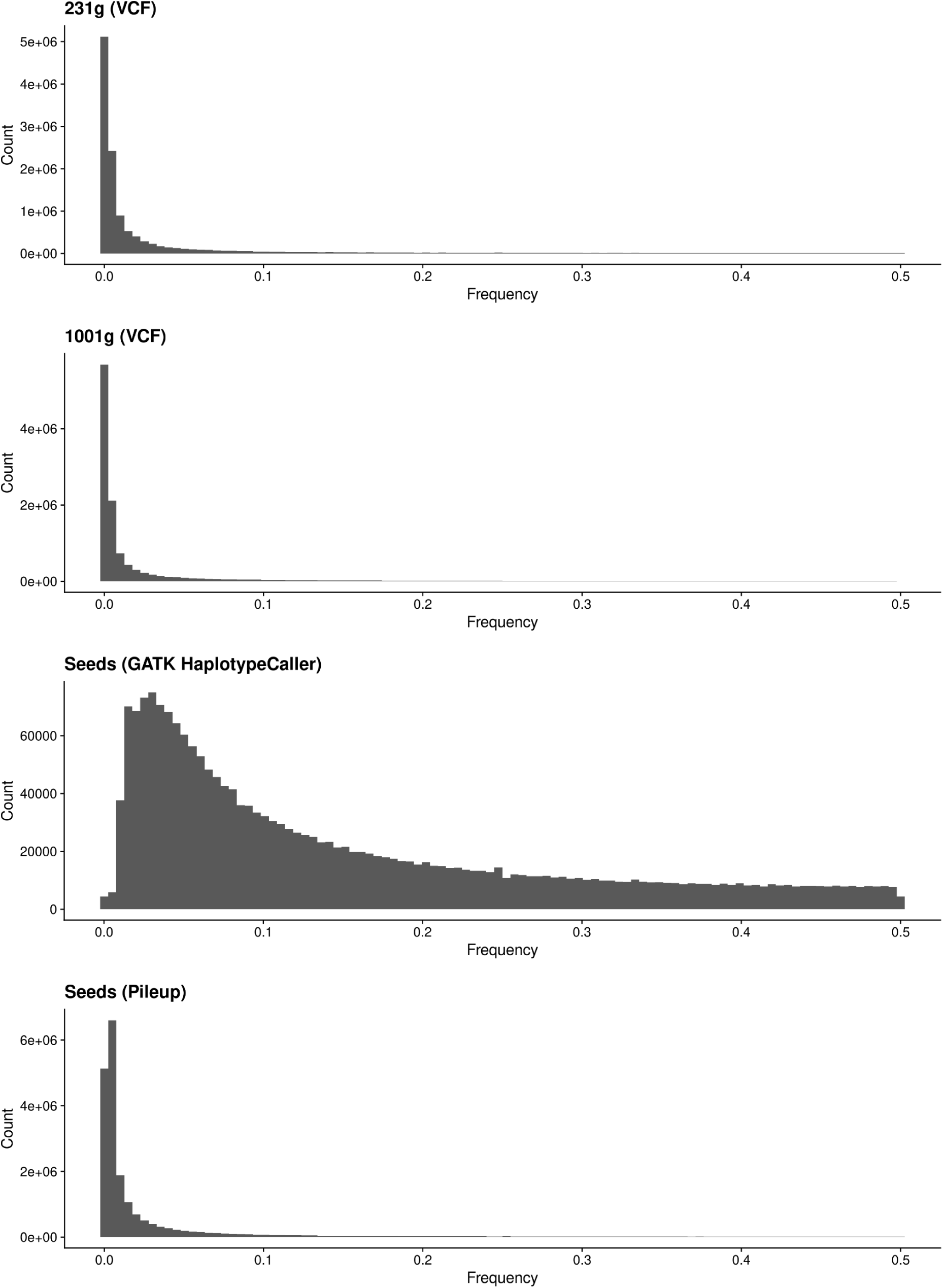
The problem of SNP calling for Pool-Seq using diploid callers. The figures show that the frequency of variants in the 1001 genomes or the 231 genomes subset is negative exponential, as expected from the Site Frequency Spectrum. The frequency calling of seeds, with high coverage, appears to be biased for intermediate frequencies in GATK HaplotypeCaller but retains the exponential decay using ratios of bases based on pileup using g_r_enedalf in the last panel.

**Fig. S11.**
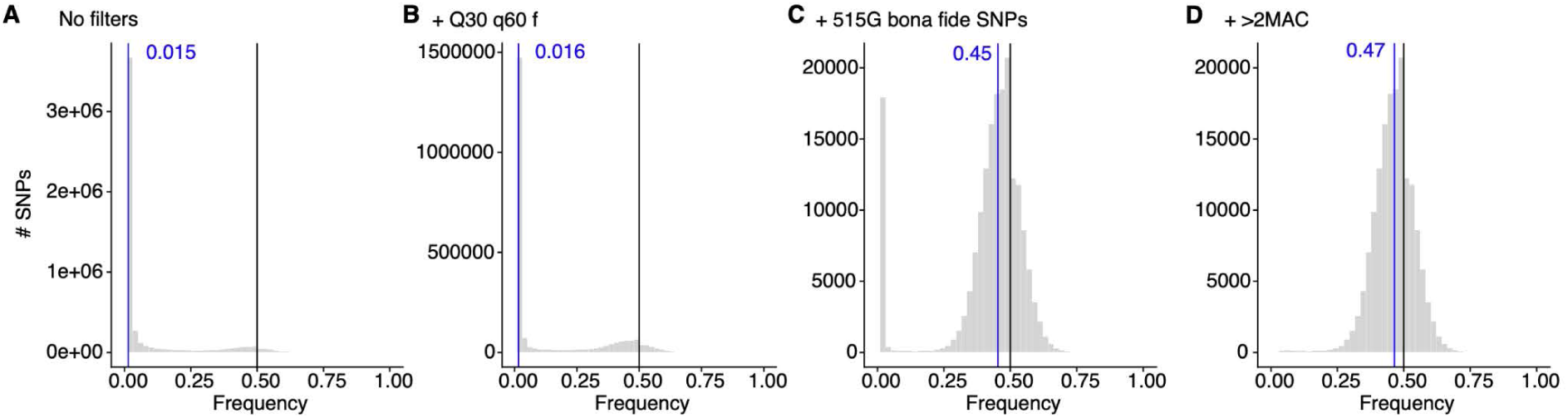
Allele frequencies in the 2 equal mass DNA pool using different quality filters. Histograms of allele frequencies from a pooled library of 2 distinct ecotypes and the expectation of 50% (black line). (**A**) Allele frequencies without any filter show the great majority of alleles must be artifacts, as there is a high point mass close to 0 frequency. (**B**) Reduction of likely artifacts, yet still high noise, using quality filters of bases with PHRED score above 30 (Q30) and from reads with mapping quality over 60 (q60) and for reads where forward and reverse map to the same region (f). (**C**) Subsetting allele frequencies to only those SNP found in the 1001 Genomes (1001 Genomes Consortium, 2016) with the highest quality 1.3 Million from <u>(Exposito-Alonso et al. 2019)</u> mostly removes all the noise signal with the exception of some rare variants. (**D**) Final removal of SNPs with only 1 or 2 bases supporting the alternative allele (minimum allele count >2) finally leaves a clean Binomial distribution of allele frequencies (owed to limited coverage) around the expected 50%.

**Fig. S12.**
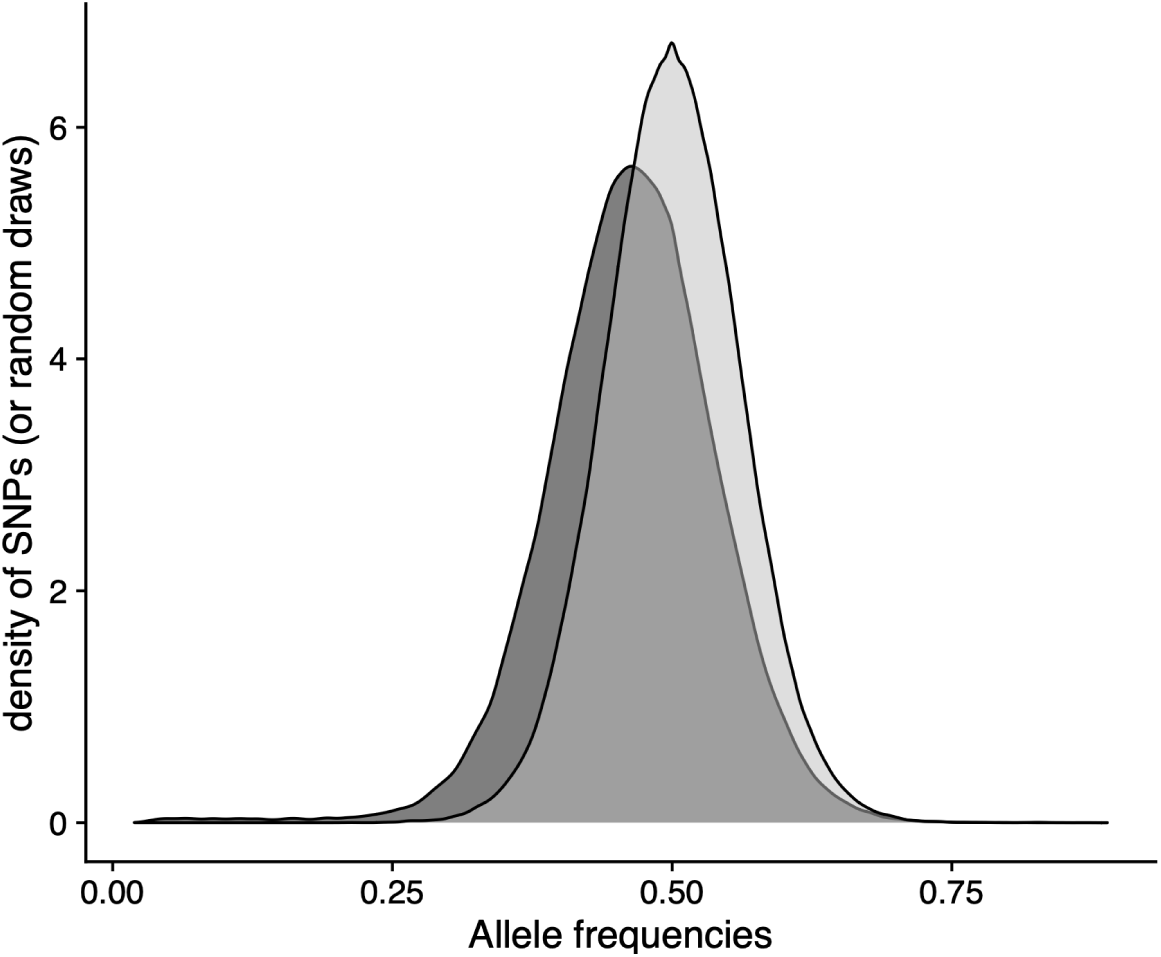
Example random Binomial draws and recovered allele frequencies. The distribution from **Fig. S12D** is compared to a random Binomial distribution with expected average frequency 50% (gray) and the same coverage distribution as the empirical sample.

**Fig. S13.**
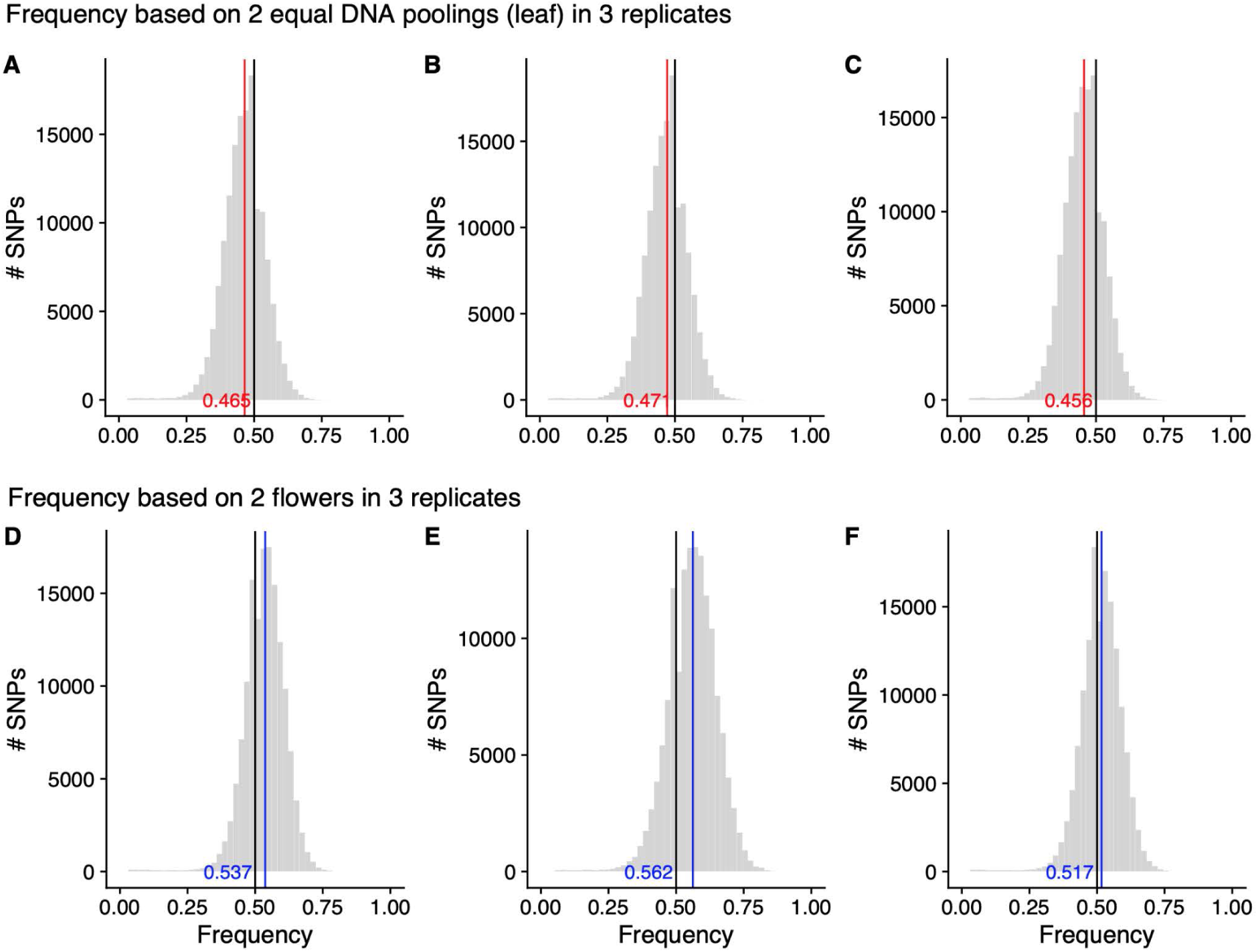
Fraction of DNA contribution to Pool-seq for a 2-flower pool and a 2-leaf pool. Dispersal of allele frequencies from the expected 50% (black vertical lines) for a pool of 2 DNA sources at equal concentration (red, **A-C**) and two flower pools (**D-F**). Both replicated three times.

**Fig. S14.**
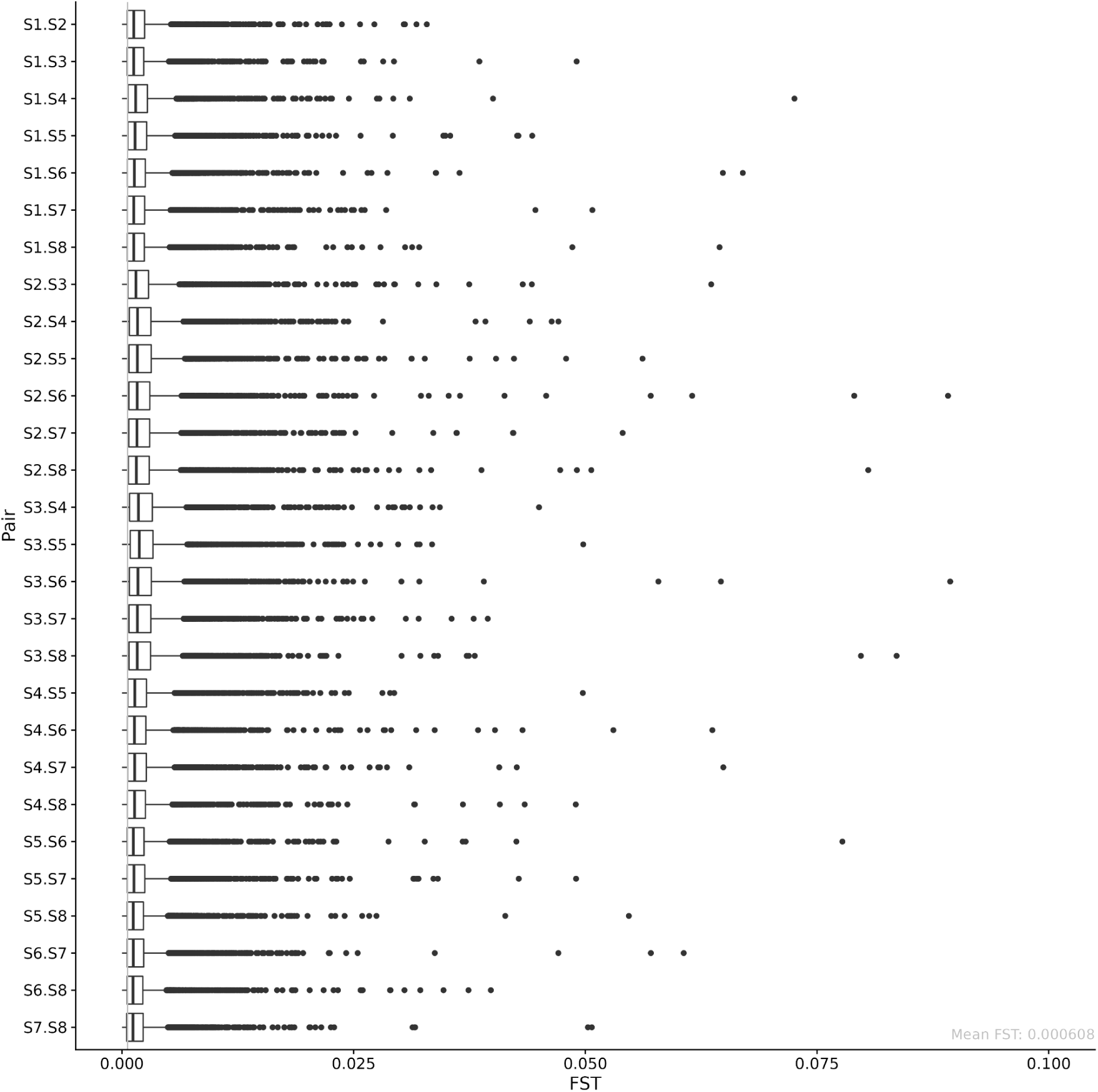
*Fst* distributions between all pairs of seed libraries. Here, we computed Fst between all pairs of sequencing of the seeds (Experiment 1) in windows of 10k base pairs across all five chromosomes, based on frequencies from the mapped data (bam/pileup files), and using the unbiased pool-seq Nei estimator as described in the main text. The window size was chosen to roughly fit the expected LD decay in *A. thaliana*. The boxes show the 25th, 50th, and 75th percentiles (i.e., quartiles), with whiskers extending to 1.5 times the interquartile range (distance between the first and third quartiles). Data points outside this range are plotted as individual points. The overall average Fst between all pairs is 0.000608, shown here as a gray vertical line, which represents the biological and statistical noise in population structure between replicates, and hence is the lower bound and baseline that we expect in other comparisons of Fst, as shown in the Figures below.

**Fig. S15.**
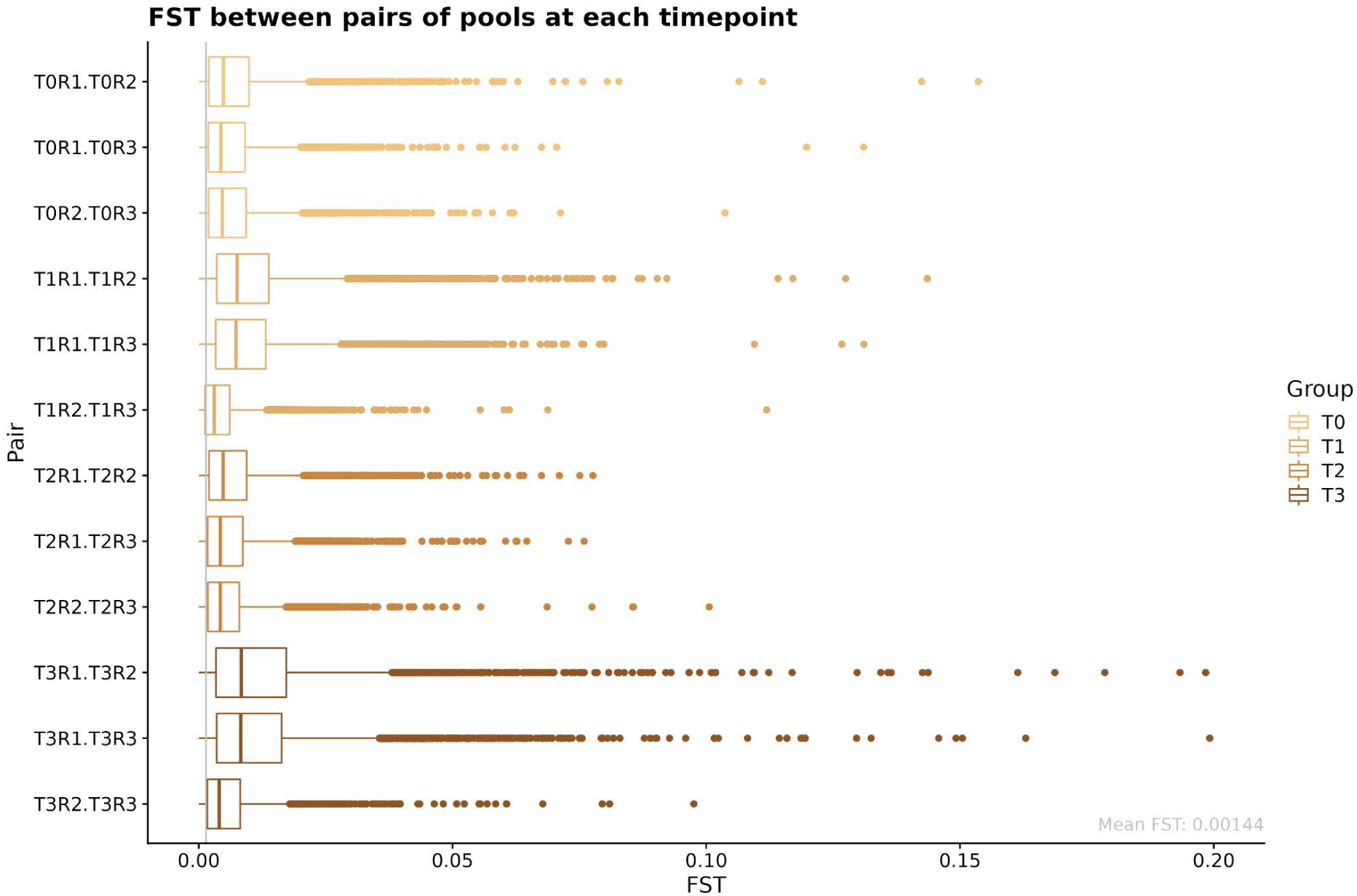
*Fst* distributions of all replicates of E&R (Exp. 4). Here, we used the data from Experiment 4, which are three technical replicates (R1-R3) grown from the same seed mix (here encoded as “time point” T0), where flowers were collected at three different time points (T1-T3) during flowering time in the spring of 2016. We here show Fst between all pairs of replicates, across all time points, in windows of 10k base pairs across all five chromosomes, and again using the unbiased pool-seq Nei estimator as described in the main text. The window size was again chosen to fit the expected LD decay in *A. thaliana*. Properties of the box plots are as above in **Figure S14**. The mean Fst, again represented as a gray vertical line here, is 0.00144, which is more than double the value of the seed baseline of 0.000608 in **Figure S14**. Note that the Fst of pairs that involve Replicate 1 is higher than that of the R2 vs R3 pairs. This is likely because R1 suffered a disturbance in the soil that could have created a bottleneck in this population and hence increased differentiation.

**Fig. S16.**
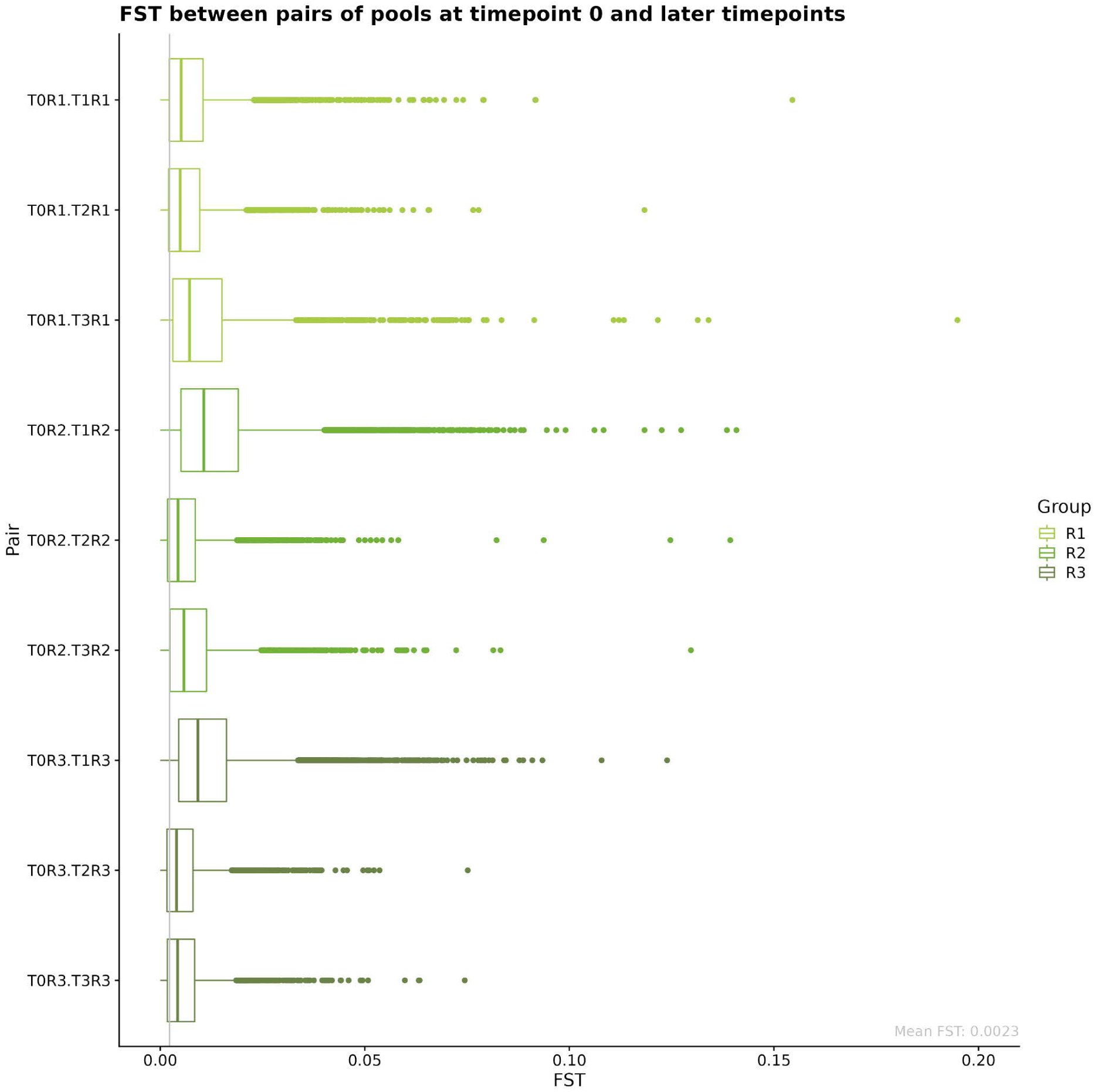
*Fst* distributions of Time 0 and all replicates in E&R (Exp. 4). Here, we used the data from Experiment 4, which are three technical replicates (R1-R3) grown from the same seed mix (here encoded as “time point” T0), where flowers were collected at three different time points (T1-T3) during flowering time in the spring of 2016. We here show Fst between the seeds and the flowering time points for each replicate, in windows of 10k base pairs across all five chromosomes, and again using the unbiased pool-seq Nei estimator. The window size was again chosen to fit the expected LD decay in *A. thaliana*. Properties of the box plots are as above in **Figure S14**. The mean Fst across all replicates and timepoints, again represented as a gray vertical line here, is 0.0023, which is almost four times the value of the seed baseline of 0.000608 in **Figure S14**. This indicates that even within one generation (from seeds to flowers), there is some differentiation happening, which suggests that rapid adaptation to the local environment of the field site has taken place.

**Fig. S17.**
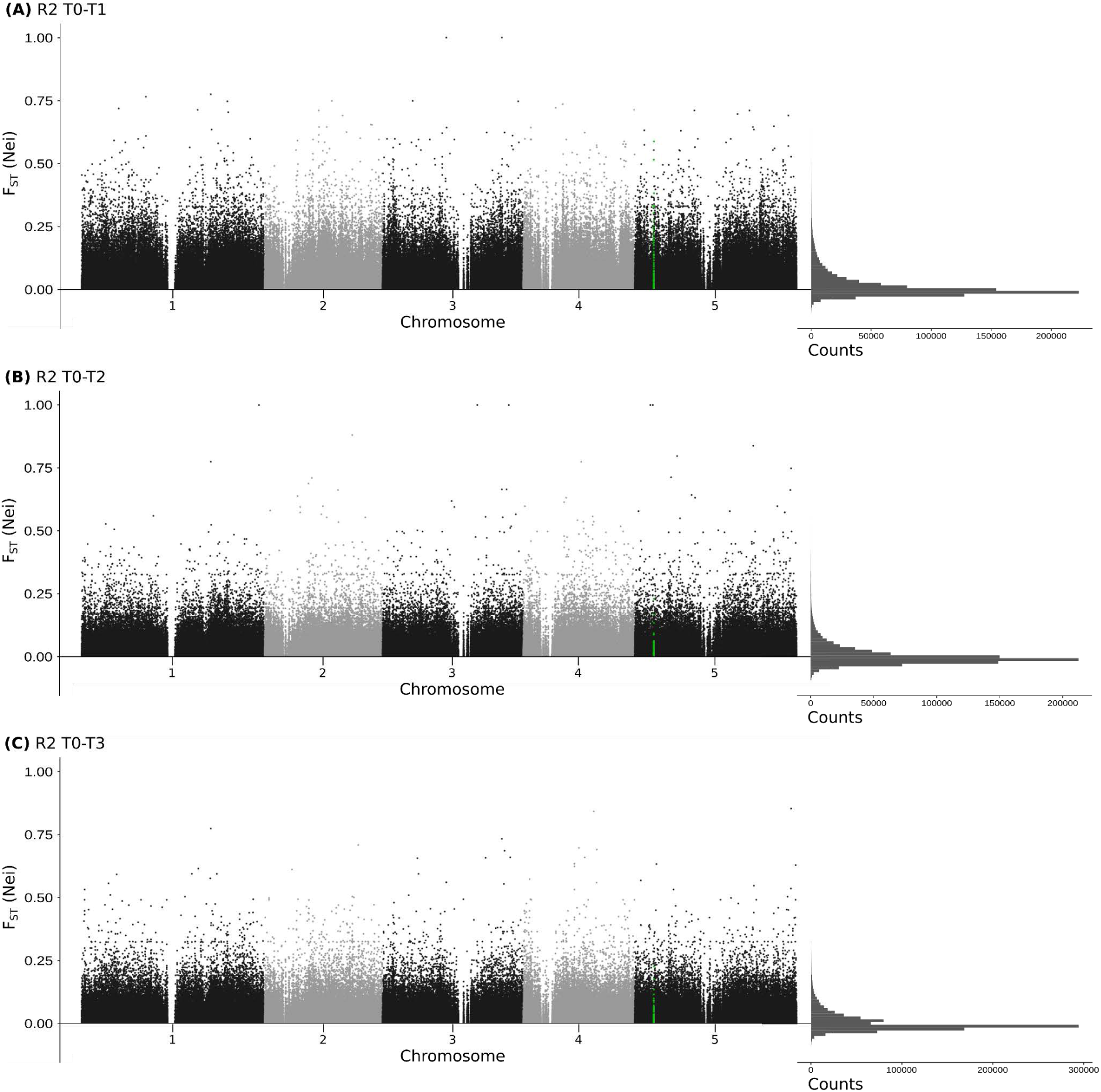
Example genome-wide *F_ST_* scan of E&R (Exp 4). Genome-wide per-SNP F_ST_ calculated using g_r_enedalf for replicate 2 between Timepoint 0 and Timepoint 1 (**A**), Timepoint 2 (**B**), and Timepoint 3 (**C**), using the unbiased pool-seq Nei estimator. SNPs overlapping with the *Flowering Locus C* (*FLC*) gene are highlighted in green (zoom in of the region in main Fig. 7). Only SNPs part of the *bona fide* 11,769,920 biallelic SNPs are shown. On the right hand side, we show histograms of the F_ST_ values displayed, indicating that most values are rather low, despite the fact that the plots on the left seemingly show an overabundance of elevated F_ST_ values. F_ST_ below zero (not shown on the left hand side) are a consequence of the estimator’s bias correction, see the **Supplemental Mathematical Appendix** for details.

**Fig. S18.**
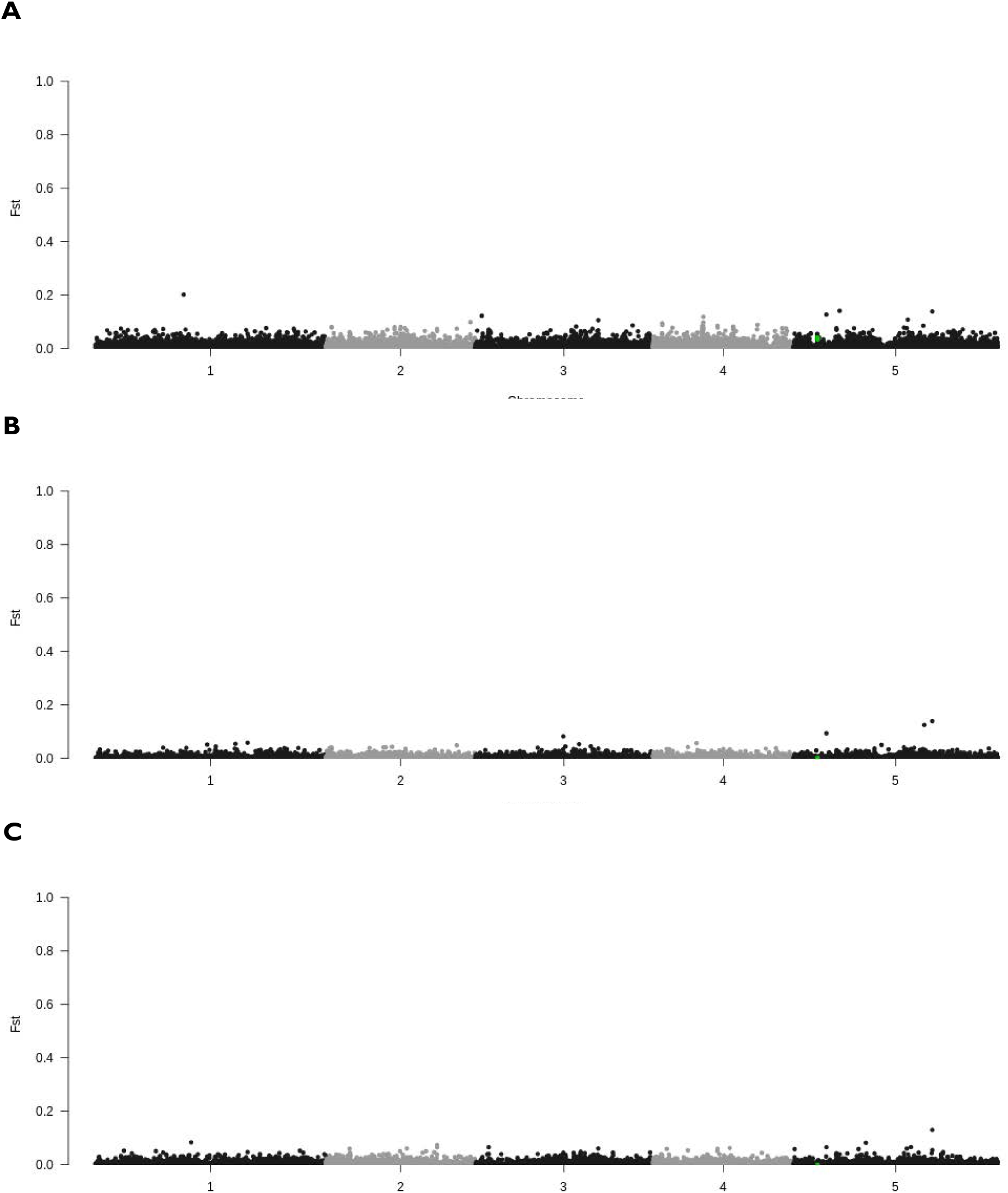
Example genome-wide *F_ST_* scan of E&R (Exp 4) with 10Kb window averages. Genome-wide *F_ST_* averages over 10Kb regions for the same samples as **Fig. S17**, again using the unbiased pool-seq Nei estimator. By computing F_ST_ in windows instead of single SNPs, most of the noise can be reduced, leaving only a few places with elevated values.

**Fig. S19.**
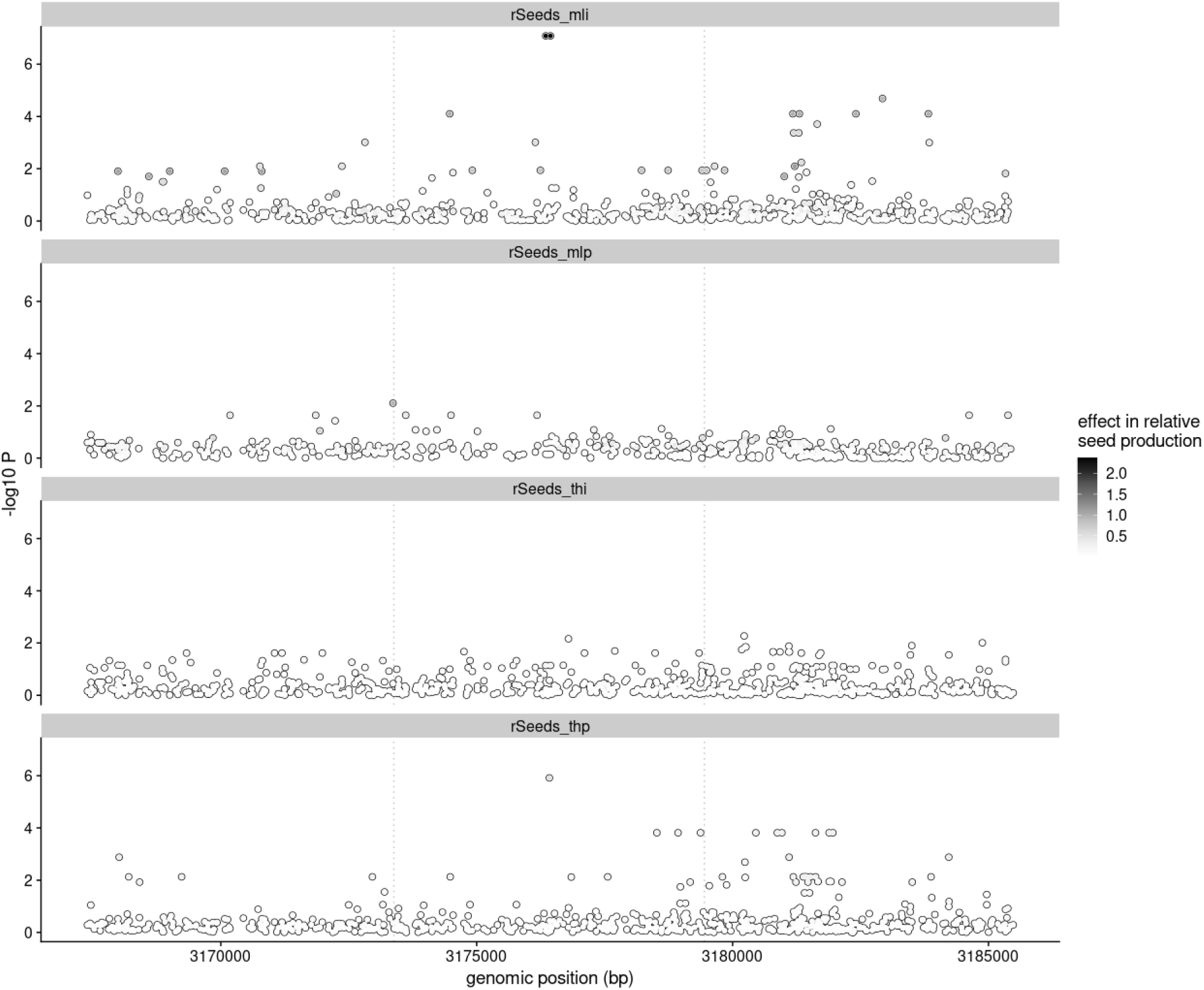
Genome-Wide Associations of different fitness component traits around FLC. Genome-Wide Association (GWA) of seed set in four key conditions from Exposito-Alonso et al. 2019. Along with condition Tübingen-high precipitation-population replicate (“thp”, Fig. 7), condition Madrid-low precipitation-individual replicate (“mli”) also show signs of SNP association to seed set within as well as the putative promoter region of the FLC gene (dotted box).

